# Exploring genome gene content and morphological analysis to test recalcitrant nodes in the animal phylogeny

**DOI:** 10.1101/2021.11.19.469253

**Authors:** Ksenia Juravel, Luis Porras, Sebastian Höhna, Davide Pisani, Gert Wörheide

## Abstract

An accurate phylogeny of animals is needed to clarify their evolution, ecology, and impact on shaping the biosphere. Although datasets of several hundred thousand amino acids are nowadays routinely used to test phylogenetic hypotheses, key deep nodes in the metazoan tree remain unresolved: the root of animals, the root of Bilateria, and the monophyly of Deuterostomia. Instead of using the standard approach of amino acid datasets, we performed analyses of newly assembled genome gene content and morphological datasets to investigate these recalcitrant nodes in the phylogeny of animals. We explored extensively the choices for assembling the genome gene content dataset and model choices of morphological analyses. Our results are robust to these choices and provide additional insights into the early evolution of animals, they are consistent with sponges as the sister group of all the other animals, the worm-like bilaterian lineage Xenacoelomorpha as the sister group of the other Bilateria, and tentatively support monophyletic Deuterostomia.

## Introduction

Large multi-gene amino acid sequence (phylogenomic) datasets promised to achieve the phylogenetic resolution [1] needed to understand the evolution of life accurately [2]. These phylogenies enable inferences about the phenotype, physiology, and ecology of common ancestors of clades [3,4], and to test hypotheses about the emergence of key innovations such as the nervous- and digestive systems [5,6].

However, modelling the evolution of amino acid sequences is difficult [7,8]. Deep metazoan phylogenies reconstructed from alternative amino acid datasets, or even the same amino acid dataset analysed using different substitution models [4,9–11], as well as using different taxon samplings of the ingroup [12,13] and the outgroup [9,10], are frequently incongruent. This acknowledged model- and data dependency of phylogenomic analyses underpins the phylogenetic instability observed towards the root of the animal tree [e.g., 14].

Although the sister group of all animals is well established – the Choanoflagellata, a group of single-celled and sometimes colonial collared and flagellated eukaryotes [15] – three nodes towards the root of the animal tree are proving difficult to resolve using multigene amino acid datasets, hindering progress in understanding early animal evolution [16].

The first recalcitrant node in the animal tree is its root, and the discussion largely centres around the question of whether sponges (Porifera) or comb jellies (Ctenophora) are the sister group of all the other animals [17,18]. This controversy impinges on our understanding of the last common ancestor of Metazoa [19], and despite receiving much attention for more than a decade [9,10,12,13,18,20–27], it is not yet resolved.

Two other recalcitrant nodes have more recently been identified from alternative analyses of amino acid datasets that affect our understanding of the root of the Bilateria (all bilaterally symmetrical animals, including humans). The first node involves the position of the worm-like Xenacoelomorpha, a bilaterian clade that unites the Acoelomorpha and Xenoturbellida [28]. With a few exceptions [29], Xenacoelomorpha are millimetre-sized and primarily benthic or sediment dwelling bilaterians devoid of a true body cavity and an anus. Xenacoelomorpha has been recovered in different positions in the animal tree: as the sister group of all other bilaterian animals (Nephrozoa) [4,29], or as the sister group of the Ambulacraria (Echinodermata+Hemichordata) constituting the clade Xenambulacraria [11,30]. The second node concerns the Deuterostomia, one of the two main bilaterian lineages (“Superphyla”). Bilateria have long been split into two lineages, Protostomia (Ecdysozoa + Spiralia [Lophotrochozoa]) and Deuterostomia (traditionally: Chordata + Ambulacraria [= Hemichordata + Echinodermata]) [31], historically based on the different origins of the mouth and other features during development [32]. However, recent phylogenomic studies challenged the monophyly of Deuterostomia and recovered paraphyletic deuterostomes in conjunction with Xenambulacraria [33,34]. This combination of results, if confirmed, would have substantial implications for our understanding of the last common ancestor of all Bilateria, which might then have been a fairly large organism, with pharyngeal gill slits and other traits previously thought to represent apomorphies of Deuterostomia ([see 34] for an in-depth discussion).

Accordingly, a stable resolution of the relationships of Xenacoelomorpha with reference to the deuterostomes is key to correctly infer the condition of the last common ancestor of the Bilateria – a small and simple organism if Xenacoelomorpha are the sister group to the Nephrozoa, or a larger and much more complex organism if Xenambulacraria is correct and Deuterostomia is not monophyletic.

Considering that previous amino-acid alignment-based phylogenomic analyses showed model- and data dependency [e.g., 9,18], which therefore did not lead to conclusive results, alternative approaches might help to select between phylogenetic hypotheses. Here we use two data types, genome gene content (“gene content”) data and morphology, to evaluate alternative hypotheses of animal relationships that emerged from previous analyses of amino acid sequence data and investigate their relative consilience [35,36]. We focus on the three recalcitrant nodes mentioned above: the relative relationships of sponges and comb jellies with respect to the other animals, the relationships of Xenacoelomorpha within the Bilateria, and the monophyly of Deuterostomia.

The phylogenetic analysis of gene content data utilises genome-derived proteomes and converts the presence or absence of gene families in the genomes of the terminals into a binary data matrix [9,25,37,38]. This approach has recently been called “molecular morphology” and advocated to also be applicable at shallower systematic levels, for example to establish a higher taxonomic system in the Placozoa [39]. We constructed separate datasets for “Homogroups” (homologous gene families) and “Orthogroups” (orthologous gene families). The former include homologous proteins that are predicted to be inherited from a common ancestor and can contain orthologs, xenologs, and out-paralogs, whereas the latter contains only proteins predicted to be inherited from a common ancestor and separated by a speciation event (see Methods for details).

We assembled a large number of new gene content datasets (see Methods, Fig. 1) to extensively test the effect of different parameter combinations when identifying homogroups and orthogroups, because this crucial step remains a challenge [40,41] and may influence the outcome of the downstream phylogenetic analysis [42]. For example, state-of-the-art methods provide two parameters (the E-value [similarity] and I-value [granulation or inflation]) which have a direct impact on the inferred gene family assignment (E-value) and splitting of gene families into orthogroups (I-value). Additionally, we explored whether removing taxa, e.g., specific outgroup taxa, before or after inferring the homogroups and orthogroups has an impact on both the size and content of datasets and inferred tree topologies.

**Figure 1:**
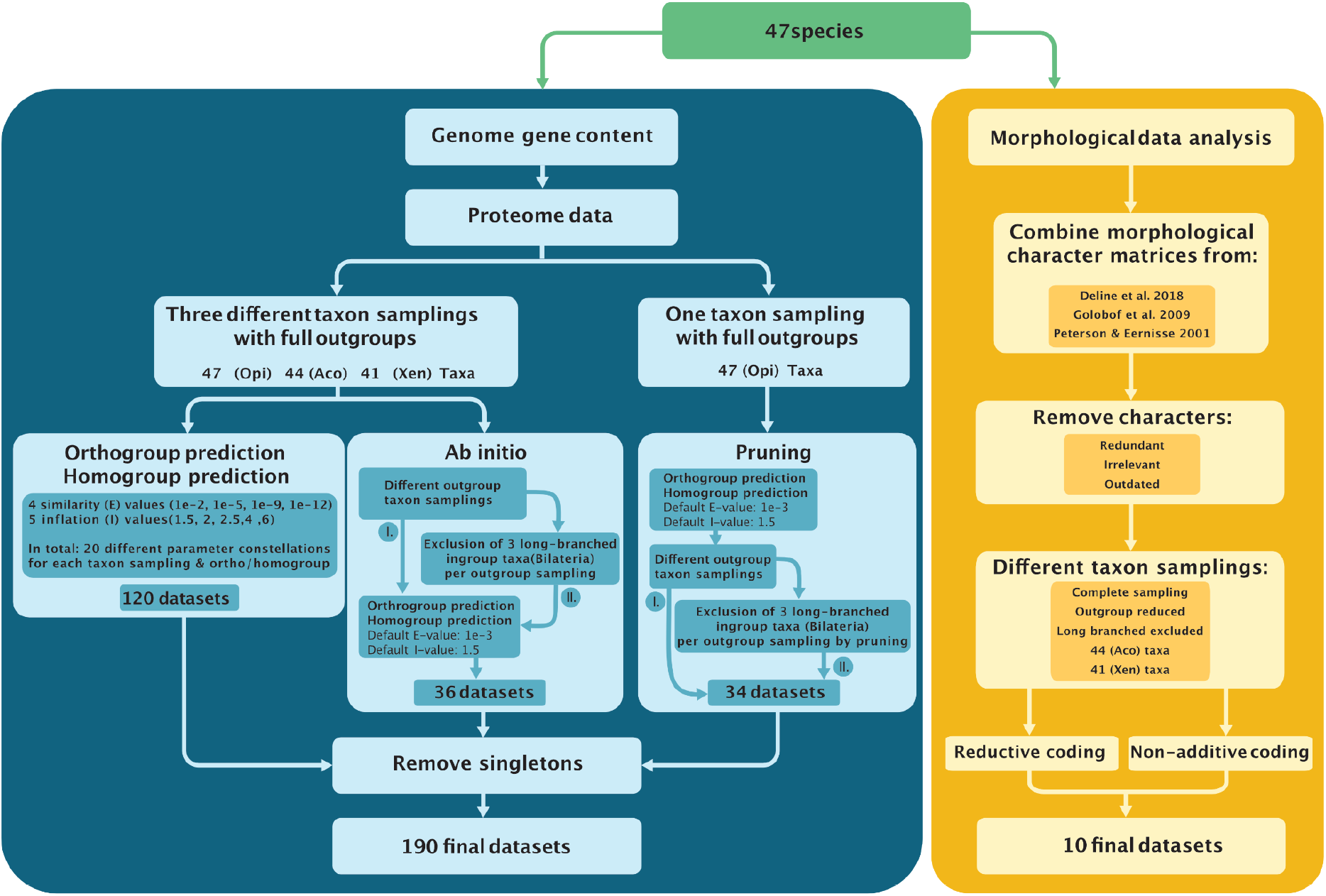
Concise graphical illustration of the methodology and workflow used for the creation of the different datasets analysed. *Left/Blue*: Genome Gene Content. “*Ab initio*” refers to dataset construction where the whole homo/orthogroup prediction was carried out *de novo* on the reduced taxon samplings, while “pruning” refers to the strategy where taxa are deleted from the full Opi homo/orthogroup data matrices which were constructed using default E (similarity) and I (inflation) values (see text for details). See <u>data repository</u> for the illustration of the complete steps of the gene content dataset creation; *Right/Yellow*: Morphology. The character list was assembled from three solid datasets that encompass the morphological disparity of the taxa in this study. Redundant characters were removed in addition to those that are not applicable to any of the terminals and historical ones that have been explicitly refuted in recent studies. The different taxon samplings mirror those of the gene content in addition to one in which the longest branches from the other morphological analyses were excluded.

We also compiled different datasets to extensively evaluate other potential sources of error, such as the so-called “long branch attraction” (LBA) artefact [43] (see Methods, Fig. 1). LBA occurs when two (or more) long branches in a phylogenetic tree group together without true relationship, generating “phylogenetic artefacts” [7]. Previous gene content analyses have focused on the root of the animals. Accordingly, here we primarily focus our LBA assessment on Xenacoelomorpha by performing taxon exclusion experiments [11]. Note that we assembled our genome gene content dataset in 2018 and included all at that time available metazoan genomes of sufficient quality. In the meantime, several new genomes have been published which could be added to our dataset. However, our study focuses explicitly on the exploration of the methods to generate genome gene content datasets from available genomes and their utility to test phylogenetic hypotheses in deep-time.

Additionally, we collated a 770-character morphological data matrix. As a starting point, we built on the classical work of Peter Ax [44] that was systematised by Deline et al. [45], and introduced additional information from two other reputable datasets [46,47] to build our matrix. All characters were reassessed before being included in our new dataset, and the coding of the base set was updated based on current morphological interpretations for groups such as Ecdysozoa and Xenacoelomorpha.

Arguably the biggest challenge when using morphology to resolve trees with very disparate taxa is character comparability, especially in our case since we have included non-metazoan outgroups. Nearly half of the characters are only applicable to either arthropods or vertebrates. The set also includes many characters that are synapomorphies of metazoan phyla, e.g., 21 characters for Nematoda. Fortunately, the base sets [45,46] contain a number of characters comparable across the whole tree. Around 20% of the characters are applicable for all metazoans, but within these some are invariable for all metazoans, or large metazoan groups. Since the outgroups are non-metazoans we also had to introduce characters specific to Fungi and Choanoflagellata [15].

In order to further mitigate the possible artefacts caused by the lack of character comparability across the tree, we utilised two different coding strategies: non-additive and reductive coding (see Methods for details). Because the non-additive coding may be affected by taxa with many uncertain states, we ran the analyses with a reduced outgroup set, which retained only the Choanoflagellata, the sister group of animals [19]. Other taxa exclusion experiments include runs without the taxa that showed problematic behaviour in the gene content analyses, the longest branches in the morphological trees, and parts of Xenacoelomorpha to check robustness. Finally, we extensively explored several modelling assumptions of morphological character evolution (e.g., ascertainment bias corrections, branch length priors, rate variation across characters and transition rate variation) to assess the robustness of our analyses. We also conducted a “total evidence” analysis, combining gene content and morphology data matrices (see Methods), to look at the contribution of the morphological data to the overall tree topology [48].

Through the provision of all analytical scripts used to assemble and analyse our datasets in a public repository (https://github.com/PalMuc/triangulation), we consider the analytical approach employed here as a template for future studies. This especially applies to the future analysis of genome gene content, once a larger taxonomic variety of chromosome-scale reference genomes becomes available, especially from undersampled non-vertebrate and non-arthropod animal lineages.

## Results

### Genome gene content data analyses

47 genome-derived proteomes were used to generate and analyse a total of 190 gene content datasets of different taxon samplings and parameter combinations (see Methods and <u>data repository</u> for details). The datasets were partitioned into several groups due to the different approaches applied (see below), all taxon sub-samplings and different parameter combinations were done in parallel for homologous gene families (“homogroups”) and orthologous gene families (“orthogroups”) [37] (Fig. 1). To assess the reproducibility of the results, the construction and analysis of the different datasets was performed twice (for results of the replicated analyses see S5 Fig; see the <u>data repository</u> for a more detailed explanation).

To test whether the specific phylogenetic relationships of Xenacoelomorpha with reference to Deuterostomia were affected by LBA, different taxon sampling experiments, based on a core taxon set of 40 species, were performed by defining three groups of datasets (Fig. 1): the “Opi” (Opisthokonta) group that consisted of all the datasets scoring a complete set of 47 taxa, including full outgroups. The “Aco” group consisted of all datasets that excluded *Xenoturbella* from the Opi dataset, and the “Xen” group consisted of all datasets that excluded the Acoelomorpha from the Opi dataset. Opi, Aco, and Xen included datasets with different parameter combinations for orthogroups and homogroups, resulting in 120 datasets in total (Fig. 1, see Methods for details).

With the same aim of detecting LBA, 70 additional datasets were generated where distant outgroups (i.e., Fungi, Ichthyosporea) and the long-branched ingroup (bilaterian) species *Caenorhabditis elegans* (Nematoda), *Pristionchus pacificus* (Nematoda), *and Schistosoma mansoni* (Platyhelminthes) were excluded, and different methods were used to construct the data matrices. Datasets were assembled using two strategies. First, the “*ab initio*” strategy carried out the whole homo/orthogroup prediction *de novo* on the reduced taxon samplings. Second, the “pruning” strategy pruned taxa from the full Opi homo/orthogroup data matrices, which were constructed using default E (similarity) and I (inflation) values (Fig. 1, see Methods for details). The *ab initio* vs. pruning dataset constructions aimed to assess the effect of those two approaches on the dimensions (gene family number) of the resulting datasets and the topology of phylogenies estimated from them.

Topologies from the individual analyses were inspected manually (see Methods, S2 and S3 Tables and S1 Fig). Additionally, Total Posterior Consensus Trees (TPCT; S4 File) were calculated for different datasets that summarise all trees sampled (after convergence) from all analyses with the exact same taxon sampling in a single majority rule consensus tree, therefore reflecting an averaging over all different E- and I-values used to reconstruct the different datasets. These trees are referred to as TPCT Opi (Fig. 2, Genome gene content), TPCT Opi-homo and Opi-ortho (S2 Fig, S5 Fig A-B), TPCT Aco-homo and Aco-ortho (S3 Fig, S5 Fig C-D), and TPCT Xen-homo and Xen-ortho (S4 Fig, Fig. 5 E-F). Support for different hypotheses was then examined using statistical hypothesis testing [49,50] (see S12 and S13 Figs).

**Figure 2:**
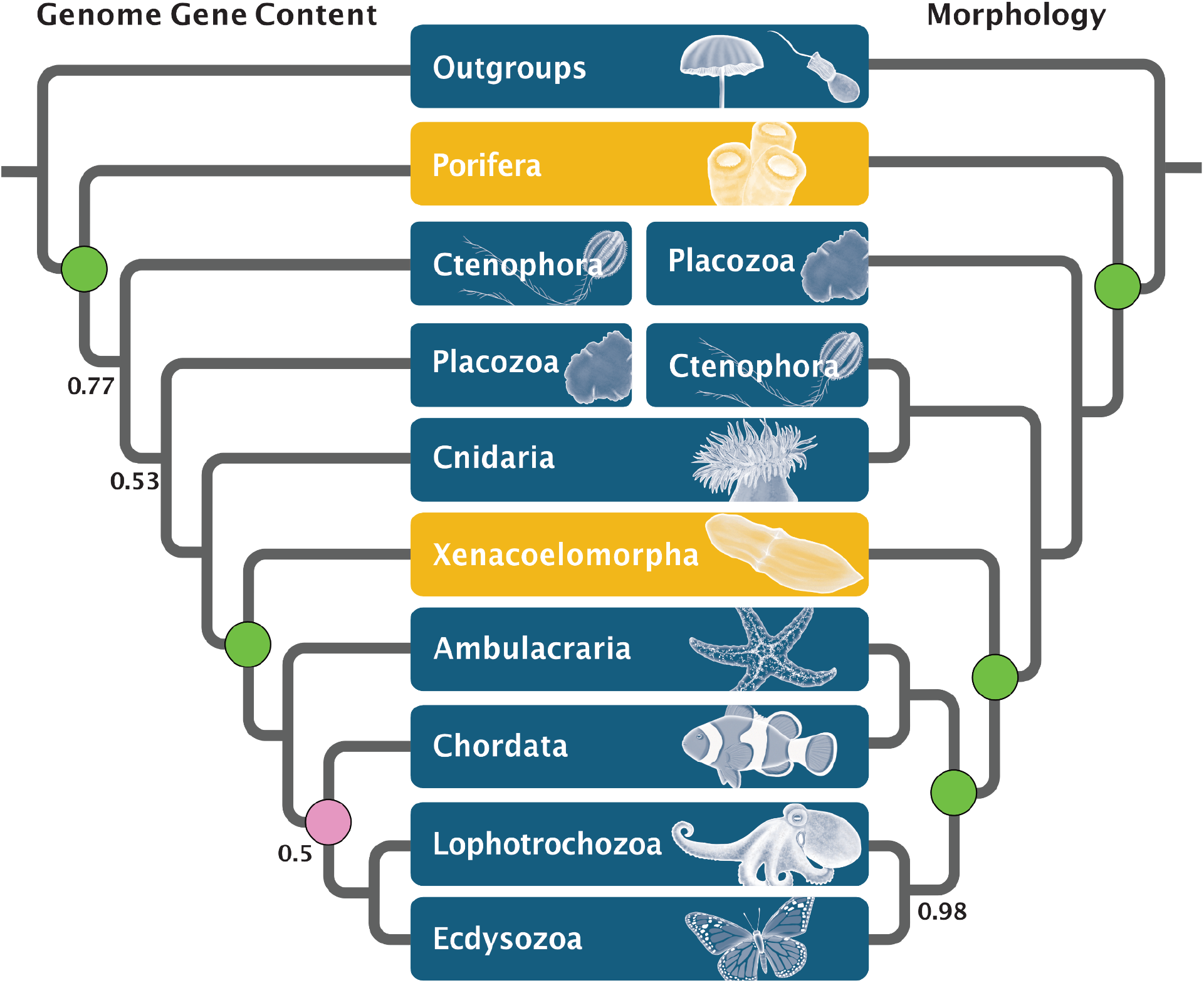
Reconstruction of animal phylogeny with 47 species (Opi taxon sampling) based on gene content datasets (TPCT) and morphological data. *Left*: Total consensus tree of >10.5 million individual tree samples from analyses using datasets of homogroups and orthogroups of all the different E- and I-values for genome gene content (for details see Materials and Methods, see S1 File for details of analytical settings). *Right*: morphology-based phylogeny based on the non-additive coding scheme. Note the different positions of Ctenophora. Second to branch off in gene content and sister group to Cnidaria in morphology (i.e., Coelenterata) analyses. The monophyly of Deuterostomia is strongly supported by morphology but around 50% by gene content datasets. Posterior probabilities lower than 0.99 are indicated on both phylogenies. Statistical hypothesis tests of focal nodes: *Green circle* = node is strongly supported in the majority of tests conducted; *Purple circle* = node is not strongly supported in the majority of tests conducted (see S9, S12, and S13 Figs for details).

### Genome gene content and the root of animals

In all 190 analyses, sponges emerged as a monophyletic group. The TPCT Opi (Fig. 2, genome gene content) indicates that the support across all analyses with a full taxon sampling is high with a Posterior Probability (PP) of 0.99 for the clade uniting all animals but the sponges, consistent with Porifera representing the sister group of the rest of the animals. Overwhelmingly strong statistical support was found for this result in our hypothesis testing (see S12 and S13 Figs; S5 Table).

Ctenophora invariably emerged as the sister group of all the animals except sponges in the TPCTs; however, the support for this node is variable in TPCTs derived from homogroups and orthogroups (PP=0.55–0.99; S2–S4 Figs). The variable level of support indicates that some analyses found Ctenophora to be placed more crownward in the tree (see Fig. 3 for the phylogeny with full taxon sampling and default I- and E-values). Three alternative topologies were found for the placement of the Ctenophora when Porifera branched first in the animals (S1 Fig C, S2–S5 Figs): Placozoa branched off before Ctenophora (e.g., Fig. 3B), the relationship between Ctenophora and Placozoa is not resolved, or Placozoa emerged as the sister group of Ctenophora. These three arrangements appear in very low numbers of trees, mostly derived from homogroup-based datasets (see S3 Table for details). In some cases, Placozoa emerges as the sister group of all animals (S3 Table). Finally, Cnidaria appears as the sister group of the Bilateria in all analyses (PP=0.99).

**Figure 3:**
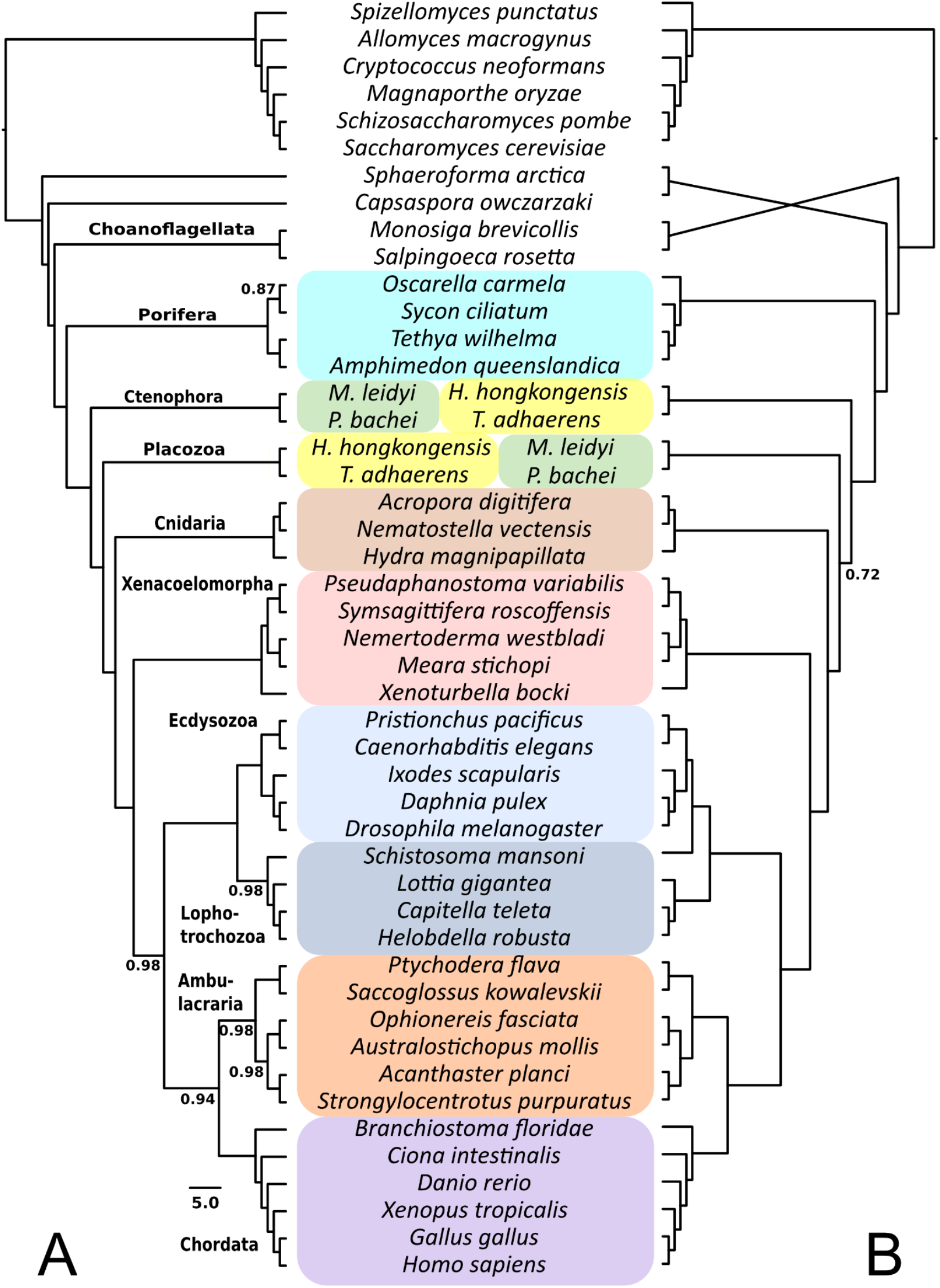
Reconstruction of animal phylogeny with 47 species (Opi taxon sampling) based on gene content datasets constructed with the default methods settings of I-value 1.5 and E-value 1e-3. Posterior probabilities lower than 0.99 are indicated. Note differences in branching order of closest outgroups, Ctenophora and Placozoa, as well as *Schistosoma mansoni* (Platyhelminthes).

### Genome gene content and the root of the Bilateria

The 47-genomes Opi dataset included five Xenacoelomorpha species and the full outgroup taxon sampling (Fig. 1, see Methods). With these datasets, Xenacoelomorpha was recovered as the highly-supported sister group of the rest of the Bilateria (Fig. 2, Genome gene content; Fig. 3), consistent with the Nephrozoa hypothesis, irrespective of whether homogroups or orthogroups were used, and with different inflation values and different outgroup sampling. Statistical hypothesis tests provided very strong support for the Nephrozoa hypothesis in 96% of the Opi, Aco and Xen datasets (S12 and S13 Figs). Similarly, datasets in the Aco group (those in which *Xenoturbella* was excluded) placed Acoelomorpha as the sister group of the rest of the Bilateria (both based on homogroups and orthogroups, S3 Fig). The overwhelming majority of the 41-genome datasets in the Xen group (those where Acoelomorpha were excluded) also resolved *X. bocki* as the sister group of the rest of the Bilateria (S3 and S5 Figs, S3 Table). Finally, in the TPCT Opi-ortho, deuterostome paraphyly is supported but with lower posterior probability (PP=0.77). Statistical hypothesis test support for deuterostome monophyly is strong from most Opi, Aco and Xen homogroup datasets, but not so from orthogroup datasets (see S12 and S13).

### Parameter changes affect mainly the final topologies from homogroup-based datasets

Different Similarity (E) and Inflation (I) values were used to construct the gene content datasets and evaluate their influence on dataset construction and downstream phylogeny estimation. Parameter changes resulted in final homo- and orthogroup matrices with different numbers of characters, but always in the range of 20,000 to 80,000 genes (S2 and S3 Tables). The choice of E-values did not significantly affect matrix reconstruction, but by contrast, the choice of I-values and whether homo-or orthogroups were used when defining matrices had significant but predictable effects.

It was expected that Orthogroup-based datasets contain a larger number of characters than the corresponding homogroup-based datasets (S1 Fig A, B), because homogroups include multiple orthogroups. Furthermore, higher inflation values resulted in the identification of a higher number of smaller homo- and orthogroups, which translated into matrices with more characters. In datasets Opi, Aco, and Xen, the lower I-values resulted in phylogenies favouring the Porifera-sister hypothesis, Xenacoelomorpha as the sister group of the Nephrozoa, and monophyletic Deuterostomia; this trend is stronger for the orthology-based datasets (see S1 Fig C).

Phylogenies based on homogroups exhibit more variability in the resulting tree topologies than phylogenies based on orthogroups. However, while the overwhelming majority of homogroup-based trees were consistent with the Porifera-sister hypothesis, 11.1% of all those trees showed Placozoa as the sister group of all the other animals. From all homogroup-based analyses that showed Porifera-sister, less than 25% of datasets constructed using high I-values placed *X. bocki* within Deuterostomia (see S1 Fig C and S3 Table). Up to 75% of homogroup-based datasets have consistent support for the Nephrozoa hypothesis, independent of inflation values.

Paraphyletic Deuterostomia appears in around 25% of the trees estimated from data sets constructed with high inflation values (S1 Fig C), while in the rest of the treatments it appears in less than 25% of the trees. The variability of the phylogenies obtained with high inflation values is also reflected in the statistical hypothesis tests performed, where high granularity of homogroups did not support any of the tested constraints (S5 File). The prediction of homo- or orthogroups appears to affect the support for deuterostome paraphyly; orthogroups favour it, while homogroup-based datasets do not (S1–S4 Figs).

The Porifera-sister hypothesis is robust to different outgroup samplings in both homogroup- and orthogroup-based phylogenies, as indicated by their very strong statistical hypothesis test support (see S5 Table). Similarly, the Nephrozoa hypothesis received very strong support from the reduced outgroup sampling datasets in our statistical hypothesis tests (see S5 Table), and all reduced taxon-sampling phylogenies where Porifera branched first supported monophyletic Deuterostomia (S1 Fig C).

The different taxon exclusion schemes showed high variations in the number of characters in the final homogroup- and orthogroup-based data matrices (S1 Fig A). However, only minor topological changes were observed in phylogenies reconstructed with different numbers of characters, compared to the phylogeny displayed in Figure 2 (Genome gene content). *Xenoturbella bocki* was only recovered in an intra-nephrozoan location in three analyses, all were from the orthogroup-based Holozoa datasets (S3 Table).

### Morphological data analyses

The morphological data sets constructed here are the first to include state-of-the-art knowledge about shared characters across Xenacoelomorpha. Two different coding schemes, i.e., non-additive and reductive coding (Methods; Fig. 1, S1 File) were applied to the morphological dataset. In addition to the different coding schemes, four taxon exclusion experiments were performed: a version with a reduced outgroup, where all the non-metazoan outgroups except the choanoflagellates were excluded from the taxon sampling, two matrices with the 41 and 44 taxon samplings (the core 40 taxa plus *Xenoturbella bocki* and the four species of Acoelomorpha, respectively) and a set without the three taxa with the longest morphological branches (dataset name Morphology Long Branches, MLB) in the previous analyses (*Ixodes scapularis* [Arthropoda], *Danio rerio, Gallus gallus* [both Chordata]). All ten analyses resulted in similar topologies (<u>see data repository for details</u>). The analysis of the non-additive matrices exhibits heterogeneous branch lengths and high node support across the phylogeny (Fig. 2, Morphology; S6 Fig). The phylogeny resulting from the datasets applying reductive coding has lower node support, with three polytomies in the ingroup (within echinoderms, chordates and the sponge classes; S7 Fig).

The only notable difference between the results of these analyses are the relationships within Porifera. In all phylogenies, sponges branched off first (Fig. 3 Morphology; S7 and S9 Fig). However, in the reductive-coding datasets, sponges are paraphyletic, with demosponges branching off first and the Homoscleromorpha and Calcarea in a polytomy with the rest of the animals. In both datasets, Placozoans branched off next and are the sister group of the traditional Eumetazoa (PP=1.0 for non-additive coding, and PP=0.89 for reductive coding). Within eumetazoans, ctenophores are the sister group of the Cnidaria (Coelenterata) (PP=1.0 for non-additive coding, and PP=0.65 for reductive coding).

In our Bayesian analyses, the hypothesis that Xenacoelomorpha is the sister group of the Nephrozoa is fully supported in the non-additive coded dataset (S 9 Fig) and the outgroup-reduced reductive coded dataset (S8 Fig), but slightly less supported in the complete sample reductive-coded phylogeny (PP=0.9) (S7 Fig). The internal relationships of Bilateria show monophyletic Nephrozoa, Deuterostomia, Protostomia, Ecdysozoa, and Spiralia in all the coding schemes applied. In order to further corroborate the results of our Bayesian analyses of the morphological data, we also analysed the set with both codings under maximum parsimony using TNT [51]. The resulting phylogenies from both codings are congruent with the corresponding results of our Bayesian analyses (S10 and S11 Figs). The differences between codings mirror the ones seen from the Bayesian analyses. The reductive coding shows paraphyletic Porifera and much lower bootstrap support overall. The only topological difference between the analyses is the support for a clade of ctenophores and cnidarians in the reductive coding. Instead of being the sister group of ctenophores, cnidarians appear in a polytomy with bilaterians and ctenophores (S11 Fig).

The statistical hypothesis tests found strong to very strong support for the topology displayed in Fig. 2 (Morphology) for the three different taxon samplings (Opi, Aco and Xen; S9 Fig). The Nephrozoa hypothesis and the Porifera-sister hypothesis have consistent very strong support. Deuterostome monophyly has strong support in the reductive coding and very strong support in the non-additive coding (see S5 Table for the exact values). This statistical support was robust over all different assumed models of morphological character evolution. However, the coding, non-additive vs. reductive, yielded different strengths of support, with the reductive coding producing weak to strong statistical support, whereas the non-additive coding produced very strong support in all scenarios (S9 Fig). Interestingly, the assumption of a fixed prior distribution over a hyperprior approach for the branch lengths reduced the strength of support in some cases (S9 Fig). None of the other modelling assumptions had any impact on the estimated strength of support for the different tested hypotheses.

### Statistical hypothesis tests tentatively support monophyletic Deuterostomia

Although the gene content TPCT displayed in Fig. 2 shows paraphyletic Deuterostomia, this tree topology received only low support (PP=0.5). Statistical hypothesis tests (S13 Fig, and details above) showed that monophyletic Deuterostomes was consistently and very strongly supported in the majority of datasets analysed, except for orthogroup taxon sampling Opi with inflation values other than the default value of 1.5, and homogroup taxon sampling Opi with higher inflation values of 4 and 6, as well as taxon sampling Xen with an inflation value of 6. The statistical hypothesis tests of the morphological data (S9 Fig) provided strong to very strong support for monophyletic Deuterostomes.

### “Total evidence” combined analysis

The combined “total evidence” phylogeny (S14 Fig) is in large parts identical to the gene content analysis from the 47-taxon Opi dataset with default I- and E-values (Figure 3). Porifera remains as the sister group to the rest of the animals, Nephrozoa and monophyletic Deuterostomia are consistently recovered. However, Placozoa and Ctenophora switch positions in the Opi-ortho+morphology dataset compared to Opi-ortho gene content-only, as Placozoa branches in the former before the Ctenophora. Another minor difference is in the position of the platyhelminth *Schistosoma mansoni* in the Opi-homo+morphology dataset compared to Opi-homo gene content-only, in the former recovered as part of the Lophotrochozoa.

## Discussion

We analysed new genome gene content datasets constructed under various settings and with various taxon samplings, and newly assembled and curated morphological character matrices. In contrast to primary sequence-based phylogenies, the use of gene content in phylogenetics is a comparably recent development [9,25,37,38] and has been advocated to complement amino acid phylogenomic analyses [14]. This approach relies on the correct estimation of the underlying ortho- and homogroups, which is affected by the tool- and parameter choices [52].

In order to gain an understanding of the effect of different parameter combinations on the prediction of ortho- and homogroups in gene content-based phylogenies, we tested a variety of similarity (E) and inflation (I) values. The differences in the numbers of characters in our datasets, as parameters change, is consistent with the observation that the identification and delimitation of gene families is difficult [40,41,53]. However, we observed good congruence across datasets over the topology in Fig. 2 (Genome Gene Content), indicating that errors induced by misidentifications of orthogroups were negligible [contra 42]), while homogroup-based topologies were less congruent mostly when high inflation values were used for the predictions.

Potential biases can be induced in the results of gene content analyses when the available genomes are fragmented, or incomplete. While we strived to use high quality genomes only, some were still fragmented, and even recent “chromosome-level” genome assemblies can not guarantee a complete and unfragmented set of the gene content of a species. For example, the genome of *Ephydatia muelleri*, not available at the time we assembled our data set in 2018, is dispersed over 1419 scaffolds, even though about 84% of it was contained in the 24 largest scaffolds, encompassing 22 of the 23 chromosomes [54]. Virtually complete chromosome scale genome assemblies of non-bilaterians are only now starting to appear, i.e., the ctenophore *Hormiphora californensis*, where 99.47% of the genome are contained in 13 scaffolds [55].

While the ascertainment bias correction introduced and used in the gene content analyses of Pisani et al. [9] and Pett et al. [37] accounts for unobserved genes in all species, no correction currently exists to account for unobserved genes in individual species, the type of bias that may be induced by incomplete genomes. However, we used ortholog and homolog identification methods that are standard in the field (see Methods) and those do not rely on complete genes, but assess the given sequence. Nonetheless, developing additional corrections to account for potential errors introduced during *in silico* genome assembly and annotation could be a fruitful avenue for future research.

Considerable attention was given to the investigation of putative long-branch attraction artefacts (LBA) that might have caused a placement of Xenacoelomorpha at the root of Bilateria and the sponges at the root of the animals. To achieve this goal we performed taxon exclusion experiments, similar to Pisani et al. [9] and Philippe et al. [11]. Based on our tests, where we do not see taxa changing position as the ingroup and the outgroup are subsampled, we suggest that the placement of Porifera and Xenoacoelomorpha in our trees does not seem to be affected by LBA.

Based on multi-gene alignments, several studies showed that the evolutionary model used can affect the inferred topologies [10,22,23,26,27,30]. For the burgeoning field of the phylogenetic analysis of gene content data, model development is still limited. Pett et al. [37] applied both the Dollo model, in which, if applied to gene content data, each gene family may be gained only once on a tree, and a reversible binary substitution model, in which a gene family may be gained more than once on a tree. Both models recovered identical topologies, but the reversible binary substitution model, also used here, was shown to have the best fit for this type of data. In any case, additional and more biologically realistic evolutionary models need to be developed to analyse genome gene content data that may show better fit and adequacy.

The estimated phylogeny from the morphological dataset is fully consistent with the results from the gene content analyses concerning the placement of Porifera and Xenacoelomorpha. A notable difference concerns the position of Ctenophora, which appears as the sister group of Cnidaria, forming the classic Coelenterata [56] (Fig. 2, Morphology). Deuterostomes are recovered as monophyletic in the morphology-based phylogeny, different from their paraphyly as recovered in a few gene content analyses. The morphological matrix also includes characters that can only be scored for one or a few taxa. These characters were retained intentionally as a conservative choice to avoid decisions that may inappropriately bias our results.

When it comes to morphological characters, the evidence available is always constrained. Some nodes close to the root of the animal tree are supported by few characters due to the low comparability of many of the characters across different body plans. Additionally, some of the key characters that support nodes close to the root might be interpreted in strikingly different ways by different scholars. For example, the presence of striated rootlets and collar complexes in feeding cells is shared by animals and choanoflagellates but the homology of these cells is disputed by some [e.g., 57]. In order to account for the controversial nature of these characters, in addition to the two separate coding schemes, we also ran a few exploratory analyses after removing these two particularly controversial characters from the matrix. In spite of the low number of characters that are supposedly informative for the base of Metazoa and their close relatives, these analyses did not affect the position of the animal phyla (data not shown). These also resulted in maximum support for Porifera-sister. Ctenophora-sister was never recovered in any of the different analyses performed on the morphological dataset.

The combined “total evidence” analyses conducted on two datasets agreed with supporting Porifera-sister, but show important differences with respect to the branching order of the Placozoa and Ctenophora. Thus, the comparably small morphological dataset with 770 characters was able to overturn the much larger gene content dataset (35,455 characters for the orthogroup dataset and 32,767 characters for the homogroup dataset). This observation supports the notion that morphological characters can have a noticeable influence on deep phylogenies in a combined “total evidence” analysis [48], and we extend this here to the combined analysis of gene content data and morphology.

Our genomic and morphological results agree with each other, with previous genome content analyses [9,37], and with phylogenetic trees of amino acid datasets supporting the Nephrozoa [4,29] and Porifera-sister hypotheses [9,12,20,22,26,27,30,58]. Our results on the other hand are in disagreement with studies that identified Ctenophora as the sister of all the other animals [10,13,21,23,25,59–61], and Xenambulacraria [11,28,30,34,62].

Nonetheless, irrespective of the arrangement of the lineages towards the root of the animal tree, the transition to animal multicellularity from a unicellular last common ancestor was marked by an expansion of a preexisting genetic toolkit to enable multicellularity [63]. The functionalities necessary for this transition, such as cell adhesion, were already present in the closest protist relatives of animals, the Choanoflagellata [19]. Additionally, new protein domains evolved in the Urmetazoan that enabled more complex traits [5,64–66], for example novel signalling pathways, such as tyrosine kinases signal transduction cascades [65] and many components of *Wnt* pathway [64], and transcription factors, such as the common glutamate GABA-like receptors [67].

Our results using an alternative approach to traditional phylogenetic analyses using amino acid data matrices showed high support for the phylogenetic hypothesis about the early animal evolution displayed in Figure 2. If we accept that sponges are the sister group of the rest of the animals (Fig. 2), it can not be excluded that the last common animal ancestor (the urmetazoan) may have been a sponge-like organism that fed using choanocyte-type cells [68]. However, the homology of the collar apparatus in the Choanoflagellata, the sister group of animals, with the one of the choanocyte in sponges is disputed [57,69,70]. In spite of that, whatever the true phenotype and metabolic capacities [71] of this urmetazoan were, the key innovations required for animal multicellularity must have happened along the stem lineage towards this urmetazoan. Furthermore, if the Porifera-sister hypothesis is correct, the last common ancestor of animals might have lacked most recognizable metazoan cell types and organ systems, despite having the capacity to transit between different cell states similar to stem cells [69].

If we accept that Xenacoelomorpha is the sister group of the rest of the Bilateria (Nephrozoa) and Deuterostomia is monophyletic, the urbilaterian (the last common ancestor of Bilateria) might have been an acoelomate worm [4]. This contrasts scenarios [72] that posit a very complex urbilaterian that could have possessed a coelom, metameric segmentation, and many other bilaterian organ systems. The most notable feature of the urbilaterian would be the lack of any ultrafiltration organs or cell types [4,73]. This lack has been argued to be primary because most xenacoelomorphs are predators and a system for nitrogen excretion is very beneficial for animals with protein-rich diets [6]. Other notable aspects would be the presence of a blind stomach without an anus and their simple gonads which would have been more similar to those of most non-bilaterians. Nevertheless, the high morphological disparity present within extant xenacoelomorphs introduces some uncertainty about the plesiomorphic status of many features. Their nervous systems, for example, are extremely varied [74] and the presence of eyes in their last common ancestor can not be established with confidence [6].

Elucidating the origin of bilaterians is also fundamental for our understanding of the early history of our biosphere. The precise sequence of character acquisition is important because it can be correlated with the appearance of more complex body plans and new metazoan ecological guilds such as burrowers and grazers. For example, in the early Cambrian fossil record, it has been postulated that the rising abundance of burrowing bilaterian animals led to the decline of the dominant Precambrian bacterial mats and an initial diversification of ecological interactions – the “agronomic revolution” [75].

Ideally, analyses of different data sources would be reconciled using either a combined analysis or a statistical framework contrasting results. Unfortunately, a combined analysis of all data types –amino acid sequence alignments, genome gene content and morphological data matrices– is currently not available in a statistical hypothesis testing framework. Developing such a method is beyond the scope of the present work.

In summary, we analysed two lines of evidence, i.e., gene content and morphological data matrices, and investigated the robustness of different parameter constellations, including taxon sampling, on the resulting phylogenies. With reference to the root of the animals, where the debate is quite mature, and many contributions from different fields exist [9,10,12,13,17,18,20–23,25–27,30,56,58–60,76], our results are consistent with the view that sponges are the sister group of all the other animals. However, resolving the exact relationships of the Ctenophora and Placozoa with respect to the Cnidaria and the Bilateria remains a future challenge.

With reference to the phylogenetic placement of the Xenoacoelomorpha, our analyses favour the Nephrozoa hypothesis. However, the debate on the placement of the Xenoacolemorpha is much less developed [4,11,28–30,34,62], with some key new hypotheses (e.g., the non-monophyly of Deuterostomia) recently emerging [33,34]. Clearly, more studies, using different datasets and methods, as well as the development of more sophisticated evolutionary models for the analysis of gene content data, are necessary to more firmly establish the relationships at the root of the Bilateria.

## Methods

### Data set creation

#### The general strategy for assembly of the genome gene content datasets

Publically available proteomes derived from full genome sequences of 47 species were collected in 2018 (S1 Table), representing 17 phyla, to create a balanced taxon sampling across animal phyla, supplementing the taxon sampling of Pett et al. [37]. The collection of proteomes also included non-metazoan outgroups sampled across Opisthokonta (Fungi + Ichthyosporea + Choanoflagellates + Metazoa; S1 File).

The core taxon set includes 40 species (bold in S1 Table), from which additional taxon samplings were created. The 47-species Opisthokonta (Opi) taxon set contained the full set of species. Two additional taxon sets (see Fig. 1; S1 and S2 File) with different taxon samplings of Xenacoelomorpha were assembled adding species to the 40-species core set: a 44-species dataset that had four Acoelomorpha species and no *Xenoturbella bocki* (specified with “Aco” in the dataset name) and a 41 species dataset that had only *X. bocki* and no Acoelomorpha (specified with “Xen” in the dataset name). The rationale behind this taxon-pruning approach was to test for long-branch attraction artefacts in the ingroup (following Philippe et al. [11]) that may impact the relationships of Xenacoelomorpha.

For each taxon sampling strategy two datasets were generated. The first coded the presence/absence of homogroups (i.e., protein families as defined by the output of the Orthofinder-1 pipeline [77]) across taxa. This coding strategy uses the shared presence of a protein family as phylogenetic evidence. The second coded the presence/absence of orthogroups. When this second coding strategy is used, individual orthogroups within each protein family are treated as individual characters. This is the same strategy introduced and justified by Pett et al. [37]

Homology searches were performed using different parameters of similarity (E-value) in DIAMOND and granulation (Inflation value; I) in the MCL algorithm. Granulation affects the cluster size, i.e., the number of the predicted clusters (orthogroups) that will be considered members of the same homogroup (i.e., of the same protein family). Small I-values indicate coarse-grained clustering resulting in larger clusters (i.e., larger protein families with many paralogs, i.e., orthogroups). Large I-values will lead to fine-grained clustering, chopping bigger clusters into smaller ones, including fewer paralogs (i.e., fewer orthogroups) [78]. Increasing the inflation value (I) therefore leads to homogroup-based datasets with more characters.

For all species in the dataset where only coding sequences (CDS) were available, transdecoder [79] was used to extract the best possible prediction of open reading frames (ORF) and corresponding proteins. All proteomes were analysed using a general approach similar to Pett et al. [37], but with different tools. A homology search of the individual proteomes against each other was conducted with a combination of four different E-values. The search was performed using Diamond v0.9.22.123 [80] for the E-values of 1e-2, 1e-5, 1e-9, and 1e-12. To obtain orthogroups, we used OrthoFinder v2.3.7 [81] with the Diamond option. To establish the homogroup datasets, we used homomcl [37] with a Diamond search. MCL v14-137 [78] was used to cluster the different gene sets with five I parameters: 1.5 (default), 2, 2.5, 4, and 6 [82,83]. Similar to Pett et al. [37], we applied a correction for the ascertainment bias in our phylogenetic model and removed all singletons (sequences that appear to be present in only one genome) from each presence/absence matrix (gene groups represented by single species). Both homogroup and orthogroup datasets therefore do not contain any single species homo-or orthogroups (singletons), i.e., proteins need to be shared by at least two species and at most all but two species. The final matrices of homogroup/orthogroup presence/absence for phylogenetic analyses were generated with custom python and BASH scripts. For the dataset naming convention used here, see S4 Table.

All steps of the analysis (dataset construction, phylogenetic analyses) were performed twice to ensure reproducibility, resulting in a total of 380 different datasets analysed.

#### Datasets to test for long-branch attraction artefacts (LBA)

Using the default E-value of 1e-3 and I-value of 1.5 in OrthoFinder, Diamond, and MCL, we further tested the outcome of different species combinations. The complete taxon sampling of the 47 Opisthokonta (Opi) species and the two subsets Aco and Xeno were used to construct further reduced datasets for two different approaches (see Fig. 2). These are divided into two sub-categories to test for putative long-branch attraction artefacts by either outgroup taxa exclusion or by excluding long-branched ingroup taxa from the taxon sampling.

#### Taxa exclusion experiments

We tested the effect of reducing taxa in two different ways: first we excluded taxa before running homology searches. When this approach is used, taxa are excluded before the datasets are generated, this is the *ab initio* approach (see Fig. 1 and S1 File). The second approach, here called “pruning” (see Fig. 1 and S1 File) simply removed taxa from the datasets. The latter significantly reduces computational time.

##### 1. Outgroup taxon exclusion

i. All outgroups but the Choanoflagellates, the sister group of the Metazoa, were successively excluded from the full 47-species Opisthokonta (Opi) taxon set, and a new OrthoFinder search was conducted to create three different taxon sets, namely ii) Ichthyosporea + Choanoflagellata + Metazoa (= Holozoa; dataset prefix Holo), and iii) Choanoflagellata + Metazoa (= Choanozoa; dataset prefix Cho) [84]; see S1 File for more details.
ii. All outgroups but the Choanoflagellates were pruned from the whole taxon set above. However, the initial character matrix derived from the full Opi dataset was used (no new OrthoFinder search), deleting new singletons and orphans (that resulted from taxon deletion) instead of re-running OrthoFinder; see S1 File for more details.

##### 2. Exclusion of long-branched ingroup taxa

i. The long-branched species *Caenorhabditis elegans* (Nematoda), *Pristionchus pacificus* (Nematoda), and *Schistosoma mansoni* (Platyhelminthes) were excluded from each of the different taxon sets described above. The complete analysis of ortho- and homogroups estimation was rerun from start to end (*ab initio*). The datasets analysed were Opi-homo/ortho-Ab, Hol-homo/ortho-Ab, and Cho-homo/ortho-Ab, where Ab refers to *ab initio*; see S1 File for more details.
ii. The long-branched species *Caenorhabditis elegans* (Nematoda), *Pristionchus pacificus* (Nematoda), and *Schistosoma mansoni* (Platyhelminthes) were excluded from the final matrix of 47 species together with the outgroups, but without re-running the complete analysis of ortho- and homogroups estimation from start to end, creating three more datasets: Opi-homo/ortho-P, Hol-homo/ortho-P, and Cho-homo/ortho-P, where P refers to *pruning*; see S1 File for more details.

Overall, 70 datasets were generated combining alternative taxon sampling and character coding (homogroups and orthogroups) strategies. For a full illustrated explanation of the different datasets created, see Fig. 1 (main manuscript) and Figure “All_graph.p.pdf” of the data repository in folder “Additional information”.

#### Phylogenetic analysis based on genome gene content data matrices

All matrices were analysed with the MPI version of RevBayes v1.0.14 [85,86]. The reversible binary substitution model [87,88] was used for phylogenetic analysis, as it was found to have the best fit to gene content data in Pett et al. [37] (for details see S6 File). Each run was conducted with four replicated MCMC runs of 50,000 to 80,000 generations to achieve full convergence. Convergence of the four runs was assessed with bpcomp and tracecomp of PhyloBayes v4.1c [89]. An ESS value >300 and bpdiff values <0.3 were used as thresholds to indicate convergence.

Majority rule consensus trees were calculated with bpcomp of PhyloBayes v4.1c [89] for each dataset and i) from the individual four MCMC runs of each of the matrices that achieved convergence; ii) from all posterior trees from all converged MCMC runs of homo- and orthogroup datasets, all different E-value (similarity) and inflation value (I) constellations with the same taxon samplings. The resulting phylogeny thus represents the total majority rule consensus tree of all posterior trees / samples from all the different MCMC simulations (TPCT). For a detailed methodological explanation of Total Posterior Consensus Tree (TPCT) see S4 File. The final trees were visualised with Figtree v1.4.4 [90], all the trees were rooted with the most distant outgroup (S1 Table).

#### Phylogenetic analysis based on morphological characters

The taxon sampling of the morphological data matrix was tailored to be identical to the 47-taxon Opi gene content dataset to make the results fully comparable (see data repository). The set of 770 morphological characters is a curated combination of three different previously published datasets: 1) Dataset 1 [46] was used due to its broad eukaryotic sampling, including some fungi and non-metazoan holozoans needed for the coding of the outgroups. 2) Dataset 2 [45] represented the animal backbone as the most comprehensive and exhaustive source of general animal morphological characters. 3) Dataset 3 [47] was added because it included more up-to-date interpretations of some morphological features. Although Dataset 2 [45] is an extensive dataset, it is based on the classical work of Peter Ax from 1996 [44] and, consequently, some well-established changes in the scoring of some characters were needed. For example, characters regarding cuticles and moulting not known at the time of Ax’s work to define the Ecdysozoa [91] were coded independently for “nemathelminthes” and arthropods in the original dataset.

The final character list analysed here (S3 File) was constructed by first combining the character lists of the publications as mentioned above. Then, the combined list was manually checked, and some characters were removed based on four criteria: 1) characters that were redundant (i.e., that reference the same information); 2) characters that only make reference to the specific morphology of clades that were not included in the sample; 3) highly debated characters where the homology was uncertain and has been questioned through independent lines of research, like the homology of “articulatan” (the classical grouping of annelids and arthropods) features [91]; and 4) characters that would have to be coded as unknown for most taxa because we are coding at the species level (i.e., reproductive, developmental and molecular).

In addition to the full 47 taxa set, four taxon sampling experiments were performed by pruning taxa from the full taxon samplings similar to the gene content analyses: two datasets without the two problematic/unresolved echinoderms and a subsample of Xenacoelomorpha (only Xenoturbella and only Acoelomorpha, respectively); a dataset without long branches observed in preliminary morphological analyses (*Danio rerio, Gallus gallus, Ixodes scapularis*); and lastly a dataset excluding all outgroups except the two choanoflagellates. All morphological data matrices are available in the <u>data repository</u>.

Modelling morphological evolution by using stochastic processes is more intricate than modelling molecular sequence evolution because it cannot be assumed that the same evolutionary process is acting on all characters identically. Stochastic processes for molecular evolution have extensively been studied and extended in the last three decades but stochastic processes for morphological character evolution are only recently catching up. Therefore, we explored several recently developed stochastic processes to test for potential biases in our phylogenetic estimates due to model assumptions. All our stochastic processes are variants of the Markov *k* (Mk) model, where *k* represents the number of states for a character, to model transitions between character states [92,93]. First, we explored the impact of ascertainment bias by either assuming that invariant characters were removed (Mkv model [92,93]) or by assuming that parsimony non-informative characters (i.e., autapomorphies) were removed. We expect that this ascertainment bias primarily influences branch length estimates but not topology estimates [94,95]. Second, we explored whether assuming a fixed exponential prior distribution with a mean of 0.1 expected substitutions per site per branch or a hyperprior distribution on the branch lengths has an impact on the estimated tree topology [96]. Third, we explored whether assuming that all morphological characters are evolving according to the same shared rate or if there is rate variation that can be modelled using four quantiles of gamma distribution [97]. Finally, we explored whether the assumption that all binary characters either share equal rates of transitions or the 0 and the 1 state occur in different frequencies by using a symmetric mixture model with four or five categories [98]. We explored all possible combinations of model assumptions (2 ascertainment bias corrections x 2 branch length priors x 2 models of rate variation across characters x 3 models of transition rate variation = 24 models per dataset) for each of the 10 morphological datasets (see Fig. 1; 240 analyses in total). These analyses were run in the Bayesian phylogenetic inference software RevBayes [86] using the MPI version. We used MCMC simulations to approximate the posterior distribution and ran two replicated MCMC simulations per analysis to check for convergence. Each MCMC simulation was run for 250,000 iterations with, on average, 150 moves per iteration. Furthermore, we used the Metropolis-Coupled MCMC extension with one cold and three heated chains to improve convergence.

Some of the reductive-coded sets (Opi and Aco) had convergence issues and were therefore run for up to 10 million generations to make sure they reached adequate values. In addition to the Bayesian analyses, we run parsimony analyses in parallel for each of the different taxonomic samplings on TNT v1.5 using the New Technology search option [51] and 100 bootstrap replicates.

#### Phylogenetic analysis based on combined data matrices for the “Total Evidence” analysis

We performed a Bayesian phylogenetic analysis using a combined dataset of genome gene content data and morphological data in RevBayes [86] (Scripts and datafiles are available at https://github.com/PalMuc/triangulation/tree/main/Combined_analyses). In this combined data analysis, we created two data partitions; one containing the genome gene content data as a matrix of absence and presence (orthogroups, homogroups; constructed with the default methods settings of I-value 1.5 and E-value 1e-3) and the second partition containing the morphological data (reductive coding). We applied the exact same models for each partition as in their independent analyses, that is, a reversible binary substitution model for the genome gene content data and the symmetric F81 mixture model [98] for the binary morphological data and the Mkv model [92,93] for the 3-state morphological data. For each morphological data partition we also applied a separate 4-category rate variation across characters using a gamma distribution [97]. We chose these models as they showed the best model fit on the separate data analyses. We linked the partitioned datasets together using the same tree topology with branch lengths in expected number of gene content changes. Thus, we applied an additional rate scaler for the morphological partition. We ran four replicated MCMC analyses for 100,000 iterations with 247.5 moves on average per iteration. We checked for convergence using the R package convenience [99]. We repeated the procedure twice, once for the homogroup dataset and once for the orthogroup dataset.

#### Hypothesis testing

We used posterior odds [49,50] to test statistical support for three competing hypotheses: (1) the Porifera-sister vs Ctenophora-sister hypotheses, (2) Nephrozoa vs Xenambulacraria hypotheses, and (3) Deuterostome monophyly vs Deuterostome paraphyly. Specifically, we computed the statistical support in favour of the null model M_0_ over the alternative model M_1_. Following standard statistical practice [49], we used the log-posterior odds of larger than 1 as substantial support, larger than 3 as strong support, and larger than 5 as very strong support. For a detailed explanation of the statistical hypothesis tests carried out see S5 File.

## Code availability

All data and code necessary to reproduce results are available in a public repository https://github.com/PalMuc/triangulation.

## Acknowledgements

GW, KJ, and DP acknowledge funding from the European Union’s Horizon 2020 research and innovation programme under the Marie Skłodowska-Curie grant agreement No 764840 (ITN IGNITE). GW and LP acknowledge funding from the Deutsche Forschungsgemeinschaft (DFG) through Project FLAGSHIP to GW (Wo896/20-1), and SH through a DFG Emmy-Noether Research Group (HO6201/1-1). GW and SH acknowledge funding through the Ludwig-Maximilians-Universität Munich (LMU) Munich’s Institutional Strategy LMUexcellent within the framework of the German Excellence Initiative. We also acknowledge Julie Johnson (www.lifesciencestudios.com) for assistance with Figs. 1 & 2 illustrations. We would also like to thank René Neumaier for our High-Performance Computing system’s design, administration and support; the present work would have been impossible without his careful and detailed work. Finally, we would like to thank six reviewers and Michael Tessler for their constructive criticisms that greatly improved iterations of the present manuscript.

## Author contributions

**K.J**.: co-designed the study, assembled the gene content data sets and carried out the gene content analyses, drafted and revised the manuscript; **L.P**.: assembled the morphological dataset and carried out the morphological analysis, drafted and revised the manuscript; **S.H**.: co-supervised the study and developed code for data analyses, revised the manuscript; **D.P**.: co-supervised the study, revised the manuscript; **G.W**.: conceived, designed and supervised the study, acquired the funding and provided the infrastructure, drafted and revised the manuscript.

## Data availability

All data and code are available in a public repository https://github.com/PalMuc/triangulation.

