## Supplementary material for "Exploring genome gene content and morphological analysis to test recalcitrant nodes in the animal phylogeny": S Fig

### Content of Supplementary Data and Figures File

**All data and code necessary to reproduce results are available on GitHub in the following repository https://github.com/PalMuc/triangulation 1**

**List of supplementary tables 2**

**Supplementary Data 1 - Methodology used to construct the datasets 3**

**Gene content analysis 3**

**Morphology 3**

Character list construction 3

Additional comments about the morphological topologies 4

**Supplementary Data 2 - Analyses of the genome gene content datasets 5**

**Individual trees for each combination of parameters 5**

**Individual tree for each combination of taxa sampling 6**

**Outgroup reduction 6**

**Outgroup and long branched ingroup reduction 7**

**Supplementary results and discussion from both analyses 7**

Results and Discussion of Supplementary Figure 1-A 8

Results and Discussion of Supplementary Figure 1-B 9

Results and Discussion of Supplementary Figure 1-C 10

**Ortho- and Homogroups based trees for each combination 10**

**Supplementary Data 3 - Character list of the morphological analyses 10**

**Supplementary Data 4 - Total Posterior Consensus Tree (TPCT) 11**

**Supplementary Data 5 - Hypothesis testing 12**

**Supplementary Data 6 - RevBayes scripts parameters 14**

**References 15**

**Supplementary Figures 17**

**All data and code necessary to reproduce results are available on GitHub in the following repository https://github.com/PalMuc/triangulation**

### 1. List of supplementary tables

**Supplementary Tables 1–5 are available on GitHub under the following link:**
**https://github.com/PalMuc/triangulation/tree/main/Tables**

**Supplementary Table1:** All species, sources, and details for the proteomic data.
**Supplementary Table 2:** Summary of all datasets settings and results for run 1.
**Supplementary Table 3:** Summary of all datasets settings and results for run 2, a summary of all the details and the most probable tree for each of the 190 datasets tested, count of the support for each of the unique topologies observed by the individual posterior trees.
**Supplementary Table 4:** Naming convention description for the long branch attraction tests for ingroups and outgroup-reduced datasets for long branches.
**Supplementary Table 5:** Statistical hypothesis testing calculation results.

**Supplementary Tables 6 – 8 are available in the text below.**

### 2. Supplementary Data 1 - Methodology used to construct the datasets

#### 2.1. Gene content analysis

A Figure (All_graph.p.png/pdf) showing the full pipeline of dataset construction is available at https://github.com/PalMuc/triangulation/tree/main/Additional%20information

Data and code used to construct the datasets are available in the data repository at
https://github.com/PalMuc/triangulation/tree/main/data_matrices_gene_content
https://github.com/PalMuc/triangulation/tree/main/Code

#### 2.2. Morphology

##### 2.2.1. Character list construction

In addition to the three main morphological datasets that were combined [1–3], a few additional characters were added to improve the resolution of the outgroups: five characters common to all Fungi [4] and two synapomorphies present in both choanoflagellate taxa were included in the analysis [based on 5]. Additionally, three other important characters were noticed to be missing in the combined dataset and subsequently were added: Schizocoely (char. 108), which is known to be one of the most important synapomorphies of protostomes (at least for the taxa sampled in our set, since chaetognaths and other groups have a different mode akin to Deuterostome development) and two general characters for muscles and excretory organs (char. 55 and 106). Furthermore, three missing sponge synapomorphies were added: a general character for the presence of sclerocytes (char. 33), the presence of an osculum (char. 768) and their cellular totipotency (char. 769). A few changes were made to five characters (Supplement 4). The coding of several characters related to cilia and statoliths within Xenacoelomorpha (characters 92 and 95) were modified in order to reflect up to date interpretations based on comparative anatomy [6]. The resulting combined dataset contains 770 characters, that are mostly binary absence/presence, except for five characters that have three states.

Coding strategies

In order to evaluate the effect of different coding strategies for absent character states, two separate data matrices were produced. In the first coding strategy, all absent states are interpreted as true absence (coded as 0; also known as non-additive coding), whereas the second coding strategy includes character dependencies based on the original reductive coding in Deline et al. [3], which distinguishes between absent and non-applicable. The non-additive coding is unrealistic because it does not respect the principle of character independence [7], but it is useful for our study because the taxonomic sample in our character matrix is very disparate and a significant proportion of the character states are inapplicable for many of the taxa. The outgroups, for example, are not animals, and most of our set is composed of animal characters, therefore they are scored as unknown for the majority of characters. By simplifying the scoring we can observe the general trends without the increased uncertainty.

##### 2.2.2. Additional comments about the morphological topologies

Vertebrates appear in a polytomy with Cephalochordata and Urochordata, which is in agreement with the equivocal morphological support for the three possible topologies [8]. Resolving the internal relationships within Porifera requires better taxon sampling. The taxon sampling here was intentionally restricted to match the 47 species taxon sampling of the genome gene content dataset to make results comparable and avoid biases in unequal taxon composition. Consequently, class Hexactinellida was not represented here because there are no genomes available and it could also not be included in the gene content analyses. Phylogenomic analyses generally support a sister relationship between the Calcarea and Homoscleromorpha [9], whereas the gene content dataset supports either the phylogenomic topology or Calcarea + Demospongiae. The non-additive coded matrices support Homoscleromorpha + Demospongiae based on the presence of silica spicules (see Morphological trees in the data repository), whereas the reductive coded matrix finds sponges paraphyletic at the base of Metazoa.

These differences are likely influenced by the limitations of our morphological analyses, which are very sensitive to differences in intrinsic anatomical complexity. In the non-additive set, all the complexity of eumetazoans produces a clear distinction between Porifera and Placozoa in one clade and the rest of the animals in another clade. In contrast, the reductive coding leads to a major decrease in the percentage of characters that sponges and placozoans can be coded for, introducing significant uncertainty in the estimated tree (i.e., lower posterior probabilities). The fact that the morphological dataset includes both unicellular organisms and vertebrates creates a major challenge for morphology since non-additive coding leads to very notable branch length gradients due to differences in character complexity/availability. This same issue is the source of the significantly lower node support across the whole tree when using the reductive-coded matrix. The analysis of different versions of the morphological dataset as it was being constructed showed that the relative position of ctenophores and cnidarians is dependent on coding hypotheses, but none of those analyses ever supported ctenophora-sister.

### 3. Supplementary Data 2 - Analyses of the genome gene content datasets

#### 3.1. Individual trees for each combination of parameters

In total, 380 phylogenies were generated, divided into two replicates (Run 1, Run 2), where the whole workflow from dataset construction to phylogenetic analysis was performed starting from the 47 proteomes.

Each of the 190 datasets were divided based on two main settings.

First, three taxon subgroups were created based on the number of species in each: Opisthokonta (Opi - 47 taxa), Acoelomorpha without *Xenoturbella bocki* (Aco - 44 taxa), and *Xenoturbella bocki* without Acoelomorpha (Xen - 41 taxa). Then, four different similarity (E) and five inflation (I) values (Supp. Tab. 6) were applied to analyse each of these subgroups, resulting in 20 datasets each (60 in total).

For each combination of parameters, homolog- and ortholog-based genome gene content prediction was carried out for dataset construction, resulting in 120 phylogenies in total (3 taxon samplings x 20 parameter combination (see Supp. Table 6) x 2 homologous and orthologous gene content).

| | | | | |
|---|---|---|---|---|
| 1.5 X 1e-2 | 2 X 1e-2 | 2.5 X 1e-2 | 4 X 1e-2 | 6 X 1e-2 |
| 1.5 X 1e-5 | 2 X 1e-5 | 2.5 X 1e-5 | 4 X 1e-5 | 6 X 1e-5 |
| 1.5 X 1e-9 | 2 X 1e-9 | 2.5 X 1e-9 | 4 X 1e-9 | 6 X 1e-9 |
| 1.5 X 1e-12 | 2 X 1e-12 | 2.5 X 1e-12 | 4 X 1e-12 | 6 X 1e-12 |

**Supplementary Table 6:** The 20 different parameter combinations for each dataset tested in Opi (47 taxa), Aco (44 taxa) and Xen (41 taxa) taxon samplings.

#### 3.2. Individual tree for each combination of taxa sampling

##### 3.2.1. Outgroup reduction

Second, the default I- and E-value of 1.5 and 1e-3 (in MCL and DIAMOND) were applied as described before on the three different taxon samplings (Opi, Aco, and Xen). Additionally, the outgroup sampling was reduced for each of the three different taxon samplings (Opi, Aco, and Xen) by following the taxon sampling of: i) the complete taxon sampling; ii) Ichthyosporea + Choanoflagellata + Metazoa (= Holozoa; dataset prefix Holo), and iii) Choanoflagellata + Metazoa (= Choanozoa; dataset prefix Cho) [10] , and two methodologies for dataset creation (Pruning and Ab Initio, here referred as A and B for simplicity) were applied to generate the final matrices.

This setting results in 17 new combinations. The Opi-B dataset was not possible to generate, because the Pruning method is deleting species from the final matrix to create a new matrix with fewer species, instead of rerunning all the pipeline, and this method can only create matrices with fewer than 47 taxa (the initial full Opi [47 species] dataset consisted of 47 taxa). For each combination described in the table, homologous and orthologous based gene content prediction was carried out to construct the presence/absence data matrix used for phylogenetic analyses. This resulted in 34 phylogenies (3 taxon samplings x 6 taxon samplings of outgroups and methodology combination x 2 homologous and orthologous predicted datasets, minus the 2 combinations which were not possible to generate).

The resulting combinations are:

| Outgroup sampling and method used | | Opi (47 sp) | Aco (44 sp) | Xen (41 sp) |
|---|---|---|---|---|
| Opisthokonta (no reduced outgroup sampling) | Method A | Opi-A | OpiAco-A | OpiXen-A |
| | Method B | - | OpiAco-B | OpiXen-B |
| Holozoa | Method A | Hol-A | HolAco-A | HolXen-A |
| | Method B | Hol-B | HolAco-B | HolXen-B |
| Choanozoa | Method A | Cho-A | ChoAco-A | ChoXen-A |
| | Method B | Cho-B | ChoAco-B | ChoXen-B |

**Supplementary Table 7**: The reduced outgroup sampling dataset designations.

##### 3.2.2. Outgroup and long branched ingroup reduction

The procedure described in the previous section was repeated for datasets without the ingroup species *Caenorhabditis elegans* (Nematoda), *Pristionchus pacificus* (Nematoda), and *Schistosoma mansoni* (Platyhelminthes) per reduced outgroup dataset. These are the excluded "near" long branch species (Suffix ne) in the datasets (see Supp. Table 8).

This step resulted in 36 additional phylogenies (3 taxon samplings x 6 taxon samplings of outgroups and ingroups with the methodology combination x 2 homologous and orthologous predicted datasets).

The dataset construction and phylogenetic analyses were performed twice, each time resulting in 190 phylogenies.

| Outgroup sampling and method used | | **Opi-ne (44 sp)** | **Aco-ne (41 sp)** | **Xen-ne (38 sp)** |
|---|---|---|---|---|
| Opisthokonta (no reduced outgroup sampling) | Method A | Opi-neA | OpiAco-neA | OpiXen-neA |
| | Method B | Opi-neB | OpiAco-neB | OpiXen-neB |
| Holozoa | Method A | Hol-neA | HolAco-neA | HolXen-neA |
| | Method B | Hol-neB | HolAco-neB | HolXen-neB |
| Choanozoa | Method A | Cho-neA | ChoAco-neA | ChoXen-neA |
| | Method B | Cho-neB | ChoAco-neB | ChoXen-neB |

**Supplementary Table 8**: The reduced outgroup and ingroup sampling performed according to the dataset naming list as presented in Supplementary Table 4.

##### 3.2.3. Supplementary results and discussion from both analyses

See Suppl. Tables 2 and 3 for all details in each run and Supp. Table 3 for the trends examination in the different individual trees, at
https://github.com/PalMuc/triangulation/tree/main/Tables

The overall all topologies count and support was calculated as the percentage of individual posterior trees from the total number of trees with the same study case only for run 2 (Supp. Table 3). The graphical summary of these results is displayed in Supp. Fig. 1 (see also data repository for further details). Each part of Supp Fig. 1 is described and discussed in detail below.

##### *3.2.3.1. Results and Discussion of Supplementary Figure 1-A*

**Results:** I-values have a more significant effect than E-values on the number of predicted homo-/orthogroups (gene families).

**Discussion:** While it can be expected that many protein families are evolutionary related and evolved through processes of gene duplication, the exact composition (in terms of orthology groups included) of a protein family can be difficult to identify. This is because the similarity of very distantly related paralogs can be minimal, and at some point, as we move backward in evolutionary history, whether two protein families should be merged into a single, larger, superfamily or not becomes difficult to decide. In software such as Orthofinder, the I-values are used to decide the extent to which orthogroups should be merged into a single homogroup (i.e., into the same family). With a larger I-value (high granularity), a large number of smaller (i.e., including fewer orthogroups) gene families are identified. Smaller I-value (low granularity), leads to the inference of less gene families which however include more orthogroups. Changes in E-values, differently, influence the number of sequences retained in each cluster (be that homo- or orthogroup). The smaller the E-value needed to accept an individual sequence as a member of a cluster, the smaller the number of sequences identified to belong to each cluster and the higher the number of singletons identified. Taxon inclusivity will also change with E-values, as higher E-values might fail to identify sequences from distantly related taxa as members of a given cluster. Combining the two parameters (specific choices of E- and I-values), as expected, resulted in homogroup datasets with variable numbers of characters, and their analyses resulted in a greater diversity of inferred trees. As expected, orthogroup-based datasets included more characters (homogroups generally include muliple orthogroups), and their analyses inferred less variable trees. We suggest that this result was to be expected as orthogroups can be partitioned across different homogroups (when changing the I-value), but when homogroups are atomized in their constituent orthogroups, the same set of orthogroups should be identified. Differences in orthogroup composition are driven by E-values rather than I-values. However, this does not necessarily mean that orthogroups are more reliable markers in genome gene content studies. Homo- and orthogroups have different strengths and weaknesses. Homogroups – if too high E-values are used in the context of low granularity analyses – might cluster orthogroups that are not homologous, introducing homoplasy in the data. However, homogroups accumulate losses more slowly than orthogroups. In this way, they are expected to be less homoplastic. Further studies will be needed to understand better what coding strategy is best in genome gene content studies. It can be hypothesised that homogroups inferred using optimal I- and

E-values would be most reliable, but how to identify optimal values for these parameters need to be further investigated.

Given the current uncertainty on how best to assemble datasets for genome gene content analyses, we have here taken the approach of testing a large range of I-values, E-values, and both ortho and homogroups. Results are mostly consistent with those in Fig. 2 suggesting the pattern described to be robust.

###### *3.2.3.2. Results and Discussion of Supplementary Figure 1-B*

**Results:** Matrices with the fewest taxon ingroup number (Xen datasets) show the highest variation in the ranges of predicted numbers of gene families for each treatment (e.g., Pruning method [P], outgroup reduction [Dis], and outgroup and ingroup reduction [Ne]). The largest variation was obtained for the Xen-Holozoa datasets, followed by the homogroups predicted datasets with the Ne treatment (Reduction of ingroups, species with long branches, and outgroups; see Methods). Also, the same treatment yields the largest variation of the predicted number of characters for orthogroup datasets. Both Opi and Aco taxon samplings show the same patterns of the predicted number of characters for all the treatments. The Aco-Choanozoa dataset shows smaller ranges of the predicted number of gene families for all treatments, while for the Opi-Choanozoa dataset the ranges are smaller and without outliers for the treatments of Pruning method (P), outgroup reduction (Dis), and outgroup and ingroup reduction (Ne).

**Discussion:** The results observed suggest that larger datasets (i.e., more outgroup taxa, larger taxon sample [Opi]) result in a more stable number of gene families in the different treatments. Homogroup-based datasets do not appear to show a distinct trend as well as datasets based on smaller taxon samplings (for both homo- and orthogroups). Together with the conclusions from Supp. Fig. 1A, the combination of dataset size and the type of dataset predicted (homo-/orthogroup) is crucial for the accurate prediction of gene families. Overall, applying the Pruning method shows a more stable range of predicted gene families compared to the *Ab initio* method, but this method can only be applied to reduce the species of interest from an initial fixed set of taxa. The initial set of taxa undergoes *de novo* prediction of gene families, which is a crucial step. Therefore, the secondary reduction (Pruning) has a less significant effect on the number of singletons. Thus, an initially well-balanced and large sample of taxa is crucial.

###### 3.2.3.3. *Results and Discussion of Supplementary Figure 1-C*

**Results:** The majority of the MCMC trees supported the Porifera-sister hypothesis. Both data types support the Nephrozoa hypothesis and slightly differ in their support for the monophyly of Deuterostomia. Datasets produced with higher E-values have slightly lower support for Porifera-sister (they support Placozoa-sister) and slightly higher support for the monophyly of Deuterostomia. Regarding the datasets produced using different I-values, all but Ctenophora-second have lower support datasets constructed with high I-values. The reduction of outgroups only (Dis) and ingroups with outgroups (Ne) showed the same trends in support of the three different hypotheses, i.e., Porifera-sister, Nephrozoa, and monophyletic Deuterostomia. The different outgroup samplings agree on the same overall trend, except the Opisthokonta outgroup sampling has slightly higher support for the Nephrozoa hypothesis than the other two outgroup samplings. Comparing the Pruned datasets to the *Ab initio* created datasets, the results show a similar trend and no difference apart from slightly higher support for the monophyly of Deuterostomia for the *Ab initio* datasets.

**Discussion:** The results observed suggest that the topology is shown in Figure 2 (main text) for genome gene content is stable and does not change considerably when different outgroups are used. The primary source of difference is the parameter settings used to generate the different types of data (homogroups vs. orthogroups, high vs. low E- and I-values).

##### 3.3. Ortho- and Homogroups based trees for each combination

Supp. Figs 2–5 show the TPCTs for each combination of species (Opi, Aco and Xen) based on the different data types predicted (homogroups and orthogroups) and the different settings used as described in the sections above (Suppl. Data 1 I and Suppl. Data 2 I-II).

### 4. Supplementary Data 3 - Character list of the morphological analyses

The full character list can be found here:
https://github.com/PalMuc/triangulation/blob/main/Morphology/morpho_character_list.txt

The full character matrices can be found in the data repository at https://github.com/PalMuc/triangulation/tree/main/Morphology/data_matrices.

#### 5. Supplementary Data 4 - Total Posterior Consensus Tree (*TPCT*)

The total posterior consensus tree is similar in approach to model [11] and data [12] averaging. Our motivation is that we do not condition the specific gene content dataset. For example, we produced different gene content datasets depending on our choice of similarity (E-value) and granulation (I-value). However, we do not know what the “correct” E-value and I-value should be, and therefore average our results over all different datasets produced.

Mathematically, we want to compute the posterior probability of a phylogeny given our genome dataset summed over all E-values and I-values, for example using the gene content orthogroups dataset, which gives

$$P(Tree \mid Genome\ data) \ = \ \sum_{i}^{N}\sum_{e}^{M} P(Tree \mid gene\ content\ data_{i,e}) \times P(i) \times P(e),$$

were $P(e)$and $P(i)$ are our prior probabilities for the different E-values and I-values. Additionally, we assume that for a given E- and I-value we obtain a gene content dataset from the genome dataset with probability 1, and thus omit this probability. Furthermore, for the $N$ I-values and $M$E-values, we assume an equal prior probability of $\frac{1}{N}$ and $\frac{1}{M}$ respectively. Thus, we can simplify our posterior probability to

$$P(Tree \mid Genome\ data) \ = \ \frac{1}{N \times M}\sum_{i}^{N}\sum_{e}^{M} P(Tree \mid gene\ content\ data_{i,e}),$$

where we can see that $P(Tree \mid gene\ content\ data_{i,e})$is our standard phylogenetic posterior probability which we estimated using MCMC sampling. Therefore, we can simply combine all posterior samples for the different MCMC simulations (assuming each MCMC simulation produced the exact same number of samples) with the different orthogroup datasets to compute the total posterior probability of a phylogeny, and similarly, the total posterior consensus tree averaged over all E-values and I-values.

The TPCT was computed both for orthogroup and homogroup datasets for the datasets testing the effect of different combinations of E-value and I-value, resulting in six TPCT in total (3 outgroup samplings and 2 method combinations). The summaries of the orthogroups and homogroups were performed separately and also combined the two types of the datasets into the phylogeny in Figure 2, genome gene content.

### 6. Supplementary Data 5 - Hypothesis testing

We performed statistical hypothesis testing for three competing hypotheses: (1) Porifera-sister vs Ctenophora-sister, (2) Nephrozoa vs Xenambulacraria hypothesis, and (3) Deuterostome monophyly vs Deuterostome paraphyly. The statistical hypothesis tests provide statistical significance for the various hypotheses. The most commonly used approach in Bayesian statistics is to compute the Bayes factor (BF). BF represents the support in favor of the null model $M_0$ over the alternative model $M_1$.

$$BF(M_0, M_1) = \frac{Posterior(M_0)}{Posterior(M_1)} \div \frac{Prior(M_0)}{Prior(M_1)}$$

The BF is often computed as the log-BF instead to avoid numerical imprecision. Standard statistical practice [13], defines a log-BF of larger than 1 as substantial support, larger than 3 as strong support, and larger than 5 as decisive support. Note that [13]) defined significance thresholds for twice the ln(BF): 2ln(BF) > 2 as support, 2ln(BF) > 6 as strong support, and 2ln(BF) > 10 as very strong or decisive support, which are ultimately equivalent to our thresholds (multiplying both sites by a factor of 2). Also, note that positive values of ln(BF) are interpreted as support for the null model M0 and negative values of ln(BF) as support for the alternative model M1.

The Bayes factors have been used to test for topological hypotheses in two ways: (a) as the posterior odds ratio only and (b) as the traditional Bayes factors accounting for the prior odds. The traditional Bayes factors for testing monophyly hypotheses have been criticized as being overly supportive of clades being monophyletic because the prior heavily penalizes against monophyly [14]. For example, adding more species in a clade that is not of interest changes the prior probability of the focal clades, but most likely not the posterior probabilities. In that case, the change in prior probabilities changes the Bayes factors and thus our conclusions, without any actual additional evidence. However, adding or removing species in distant clades from the focal clades should not impact our prior belief about our specific hypothesis, and therefore posterior odds should be preferred over Bayes factors [14].

We obtained the posterior odds from the posterior samples of trees by computing the posterior probability of both null and alternative hypotheses. If the posterior probability was either 1.0 or 0.0, then we subtracted or added $\frac{1}{N}$, where N is the number of MCMC samples, respectively, to avoid problems when computing the ratio. This approach is conservative (see [15]) and limits our power to compute statistical support only to a maximal precision of $posterior\ odds\ =\ N\ -\ 1$ or $posterior\ odds\ =\ \frac{1}{N-1}$respectively.

For our first question (Porifera-sister vs Ctenophora-sister) we computed the posterior probability of the clade (Ecdysozoa + Lophotrochozoa + Chordata + Echinodermata + Hemichordata + Xenacoelomorpha + Cnidaria + Placozoa + Ctenophora) being monophyletic (null hypothesis; Porifera-sister) as well as the clade (Ecdysozoa + Lophotrochozoa + Chordata + Echinodermata + Hemichordata + Xenacoelomorpha + Cnidaria + Placozoa + Porifera) being monophyletic (alternative hypothesis; Ctenophora-sister). For our second question (Nephrozoa vs Xenambulacraria) we computed the posterior probability of the clade (Ecdysozoa + Lophotrochozoa + Chordata + Echinodermata + Hemichordata) being monophyletic (null hypothesis; Nephrozoa) as well as the clade (Xenacoelomorpha + Hemichordata + Echinodermata) being monophyletic (alternative hypothesis; Xenambulacraria). For our third question (Deuterostome monophyly vs Deuterostome paraphyly) we computed the posterior probability of the clade Deuterostome being monophyletic (null hypothesis; Deuterostome monophyly) and the clade Deuterostome being paraphyletic (alternative hypothesis; Deuterostome paraphyletic).

We computed the posterior odds separately for each E-value, I-value and MCMC replicate. The posterior probabilities were computed in RevBayes and the posterior odds and posterior odds in a custom R script available in the data repository
https://github.com/PalMuc/triangulation/tree/main/Code

#### 7. Supplementary Data 6 - RevBayes scripts parameters

The code used for phylogenetic estimation from the genome gene content data matrices in this study is available in the data repository https://github.com/PalMuc/triangulation/blob/main/Code/mcmc_gene_content_original.Rev

A full description of the different steps of the code can be found in Pett et al. [16].
The main assumptions of this gene content evolution model are that there are no absent (i.e., no invariant sites) and no singleton sites included. The gene content evolution model was a binary, continuous time reversible Markov chain with estimated stationary frequencies. Additionally, we assume that homogroups or orthogroups can evolve according to one of four discrete rate categories (modeled by the mean of the quartiles of a gamma distribution), which is identical to the among-site rate variation used in nucleotide substitution models. We performed a slight modification to the original model of Pett et al. [16] by fixing the hyperparameter of the branch length prior distribution to an expectation of 0.1, as standardly applied in phylogenetic analyses (see, for example, the program MrBayes [17]).

#### Supplementary Figures

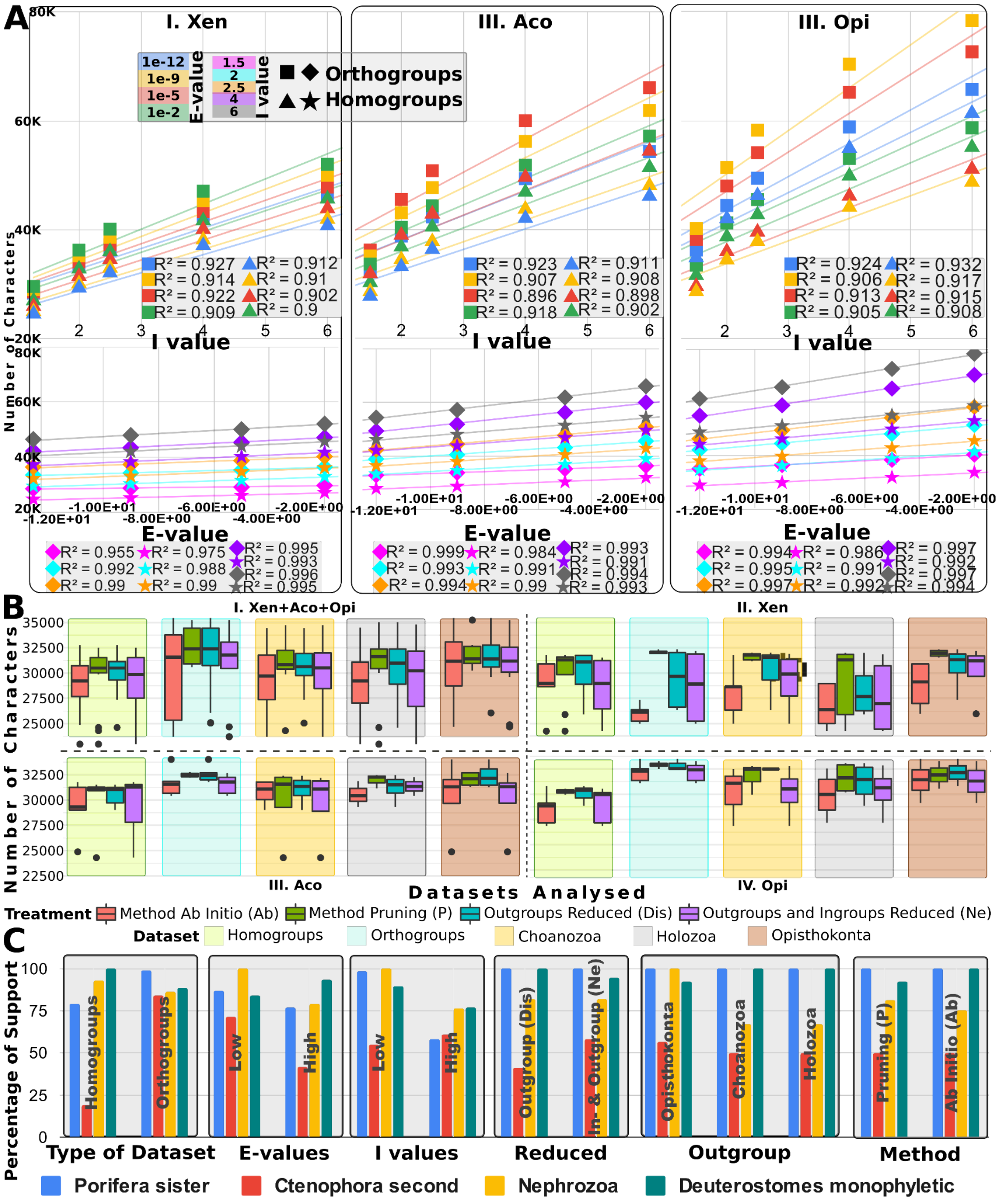


**Supplementary Figure 1: Gene content – Number of ortho-/homo- groups and main tree topology supported by each dataset and the summary of Total Posterior evidence trees.** The number of ortho-/homo- groups in each of the datasets for phylogenies represented in the scatter plots, box plots and the percentage of support for selected hypotheses are represented in the graphs. Showing the relationship between similarity (E-values) and granulation (I-values) tested on the final number of characters, as well as the outgroup choice and long branches species treatments of the dataset effects.

**A**. Numbers of characters for each combination of E-value and I-values for the three datasets Opi, Aco and Xen after the prediction of orthogroups and homogroups, trend lines and fit of $R^2$ shown (Logarithmic trend line for the I-value graphs, top and linear trend line for the E-values, bottom).
**B**. The predicted number of characters for each treatment tested in this study for all predicted datasets with E-value 1e-3 and I-value 1.5.
**C**. The percentage of individual posterior trees supporting each of the tested hypotheses in the different settings of the Opi, Aco, and Xen datasets that were used in this study. Data in Suppl. Table 3.

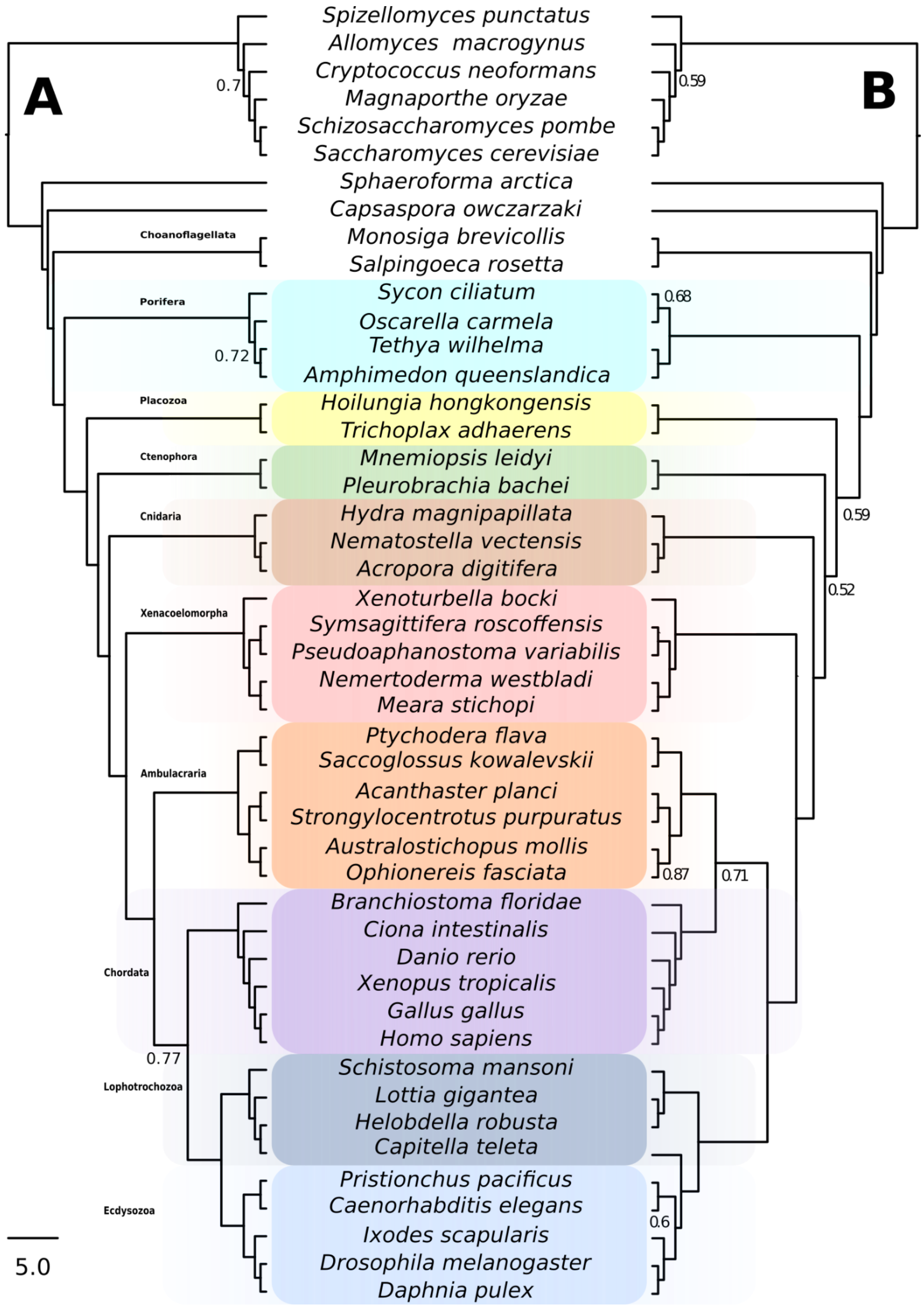


**Supplementary Figure 2**: **Gene content – Opi Orthogroups (TPCT Opi-ortho) vs Homogroups (TPCT Opi-homo).** A. Phylogeny based on orthogroups gene families predicted for 47 species. B. Phylogeny based on homogroups gene families predicted for 47 species. Each tree represents the consensus tree of 20 analyses (TCPT) performed with combinations of four E-values and five I-values. Each TPCT included samples of trees of all converged MCMC runs for each dataset analysis. The trees are presented as cladograms with proportional branch lengths. Posterior probabilities lower than 0.99 are indicated.

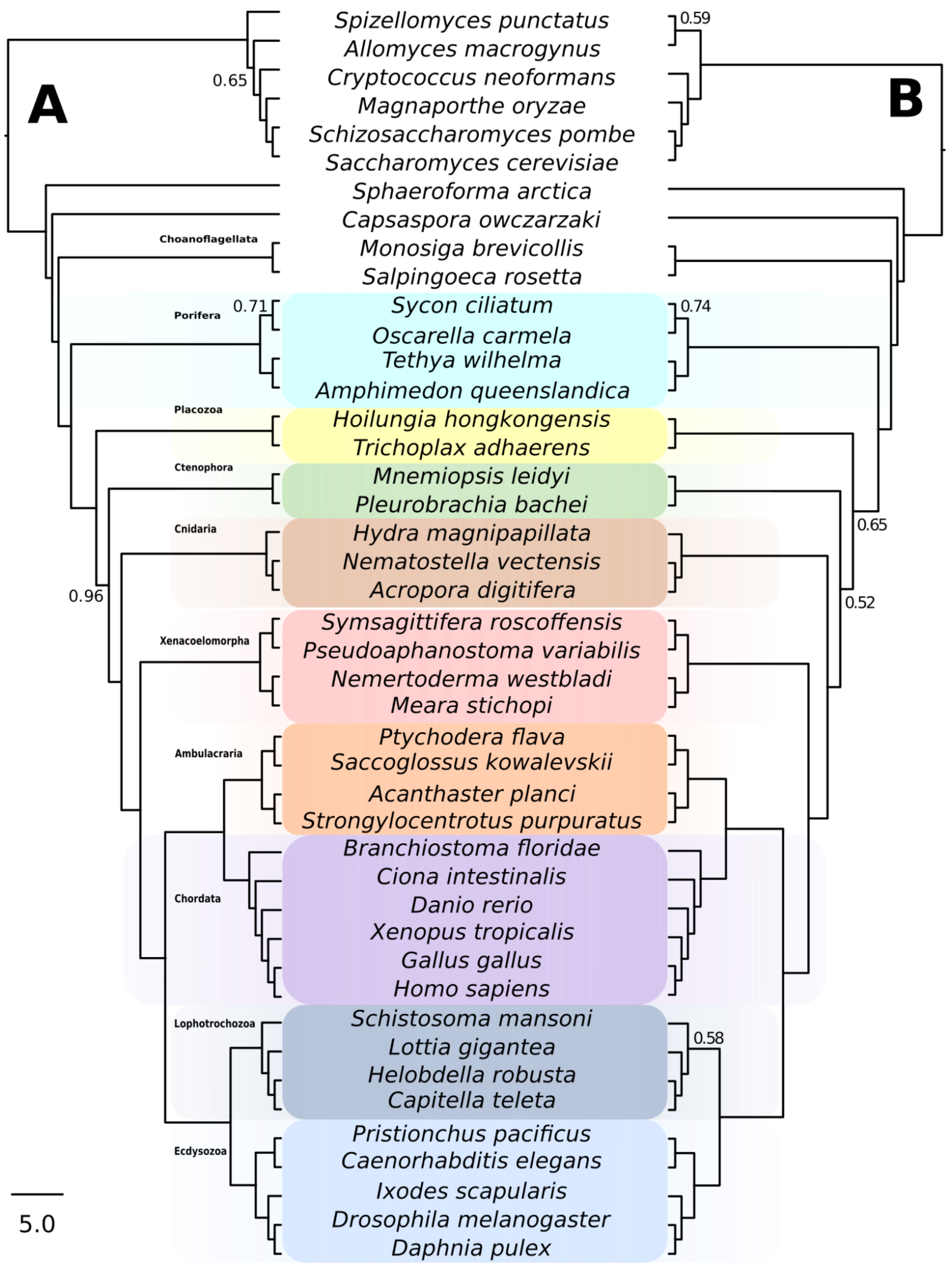


**Supplementary Figure 3**: **Gene content – Aco Orthogroups (TPCT Aco-ortho) vs Homogroups (TPCT Aco-homo).** A. Phylogeny based on orthogroups gene families predicted for 44 species. B. Phylogeny based on homogroups gene families predicted for 44 species. Each tree represents the consensus tree of 20 analyses (TCPT) performed with combinations of four E-values and five I-values. Each TPCT analysis included samples of trees of all convergent MCMC chains runs of trees for each dataset analysis. The trees are presented as cladograms with proportional branch lengths. Posterior probabilities lower than 0.99 are indicated.

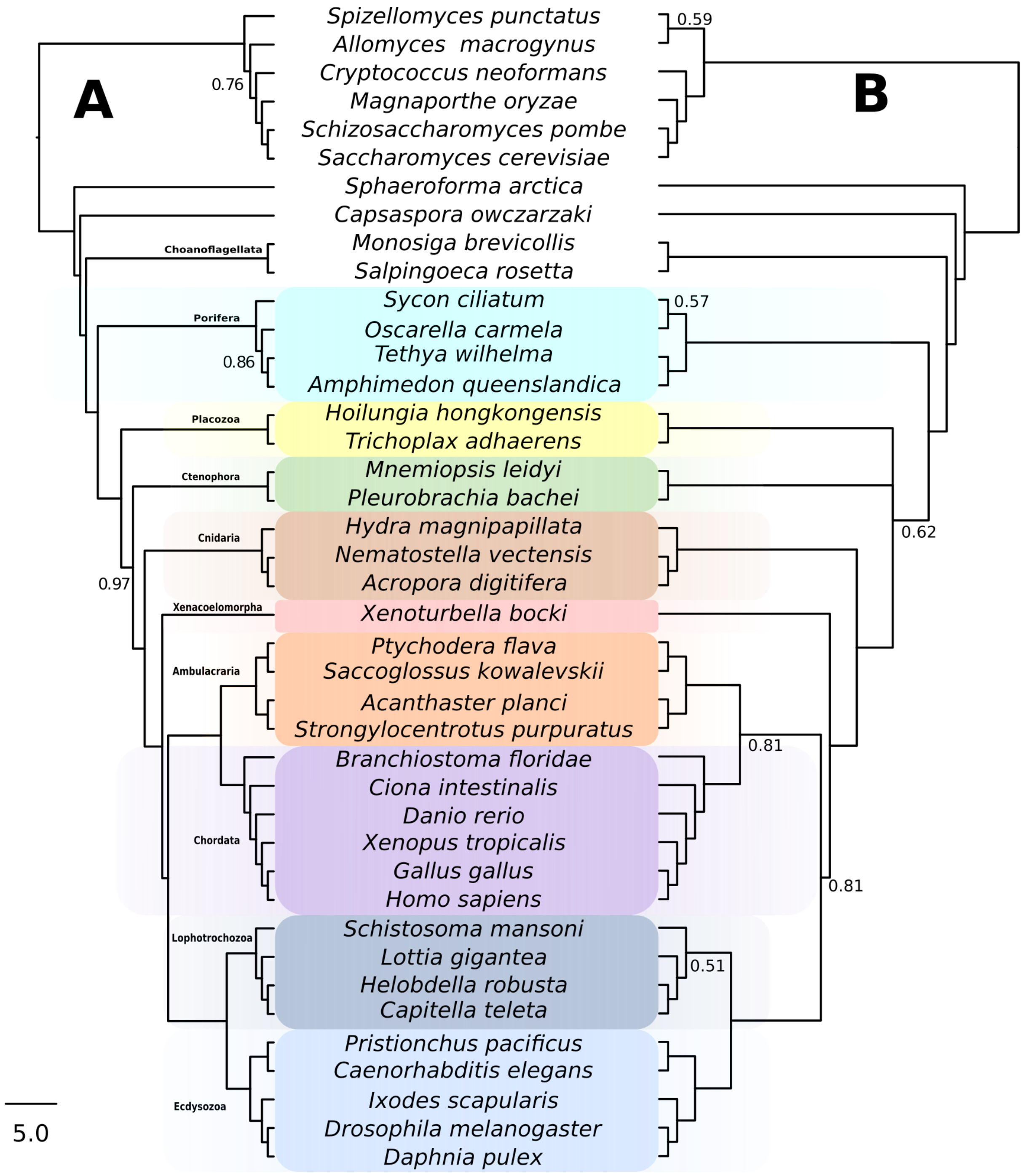


**Supplementary Figure 4**: **Gene content – Xen Orthogroups (TPCT Xen-ortho) vs Homogroups (TPCT Xen-homo).** A. Phylogeny based on orthogroups gene families predicted for 41 species. B. Phylogeny based on homogroups gene families predicted for 41 species. Each tree represents the consensus tree of 20 analyses (TCPT) performed with combinations of four E-values and five I-values. Each TPCT analysis included samples of trees of all convergent MCMC chains runs of trees for each dataset analysis. The trees are presented as cladograms with proportional branch lengths. Posterior probabilities lower than 0.99 are indicated.

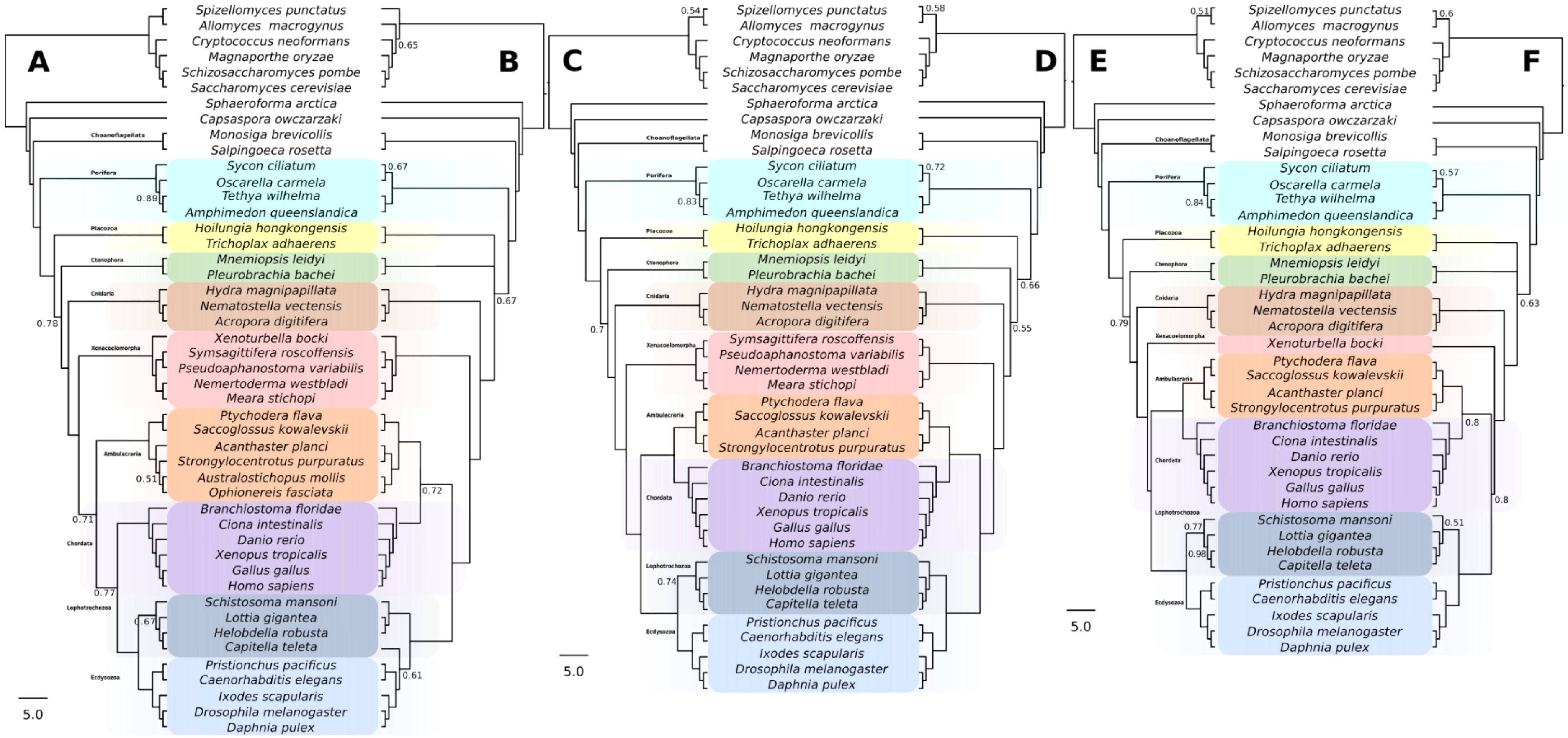


**Supplementary Figure 5**: **Gene content – TPCT trees second (replicate) run.** A. Phylogeny based on orthogroups gene families predicted for 47 species (Opi-ortho). B. Phylogeny based on homogroups gene families predicted for 47 species (Opi-homo). C. Phylogeny based on orthogroups gene families predicted for 44 species (Aco-ortho). D. Phylogeny based on homogroups gene families predicted for 44 species (Aco-homo). E. Phylogeny based on orthogroups gene families predicted for 41 species (Xen-ortho). F. Phylogeny based on homogroups gene families predicted for 41 species (Xen-homo). Each tree represents the consensus tree of 20 analyses (TCPTree) performed with combinations of four E-values and five I-values. Each tree included MCMC samples of all converged runs for each dataset. The trees are presented as cladograms with proportional branch lengths. Posterior probabilities lower than 0.99 are indicated.

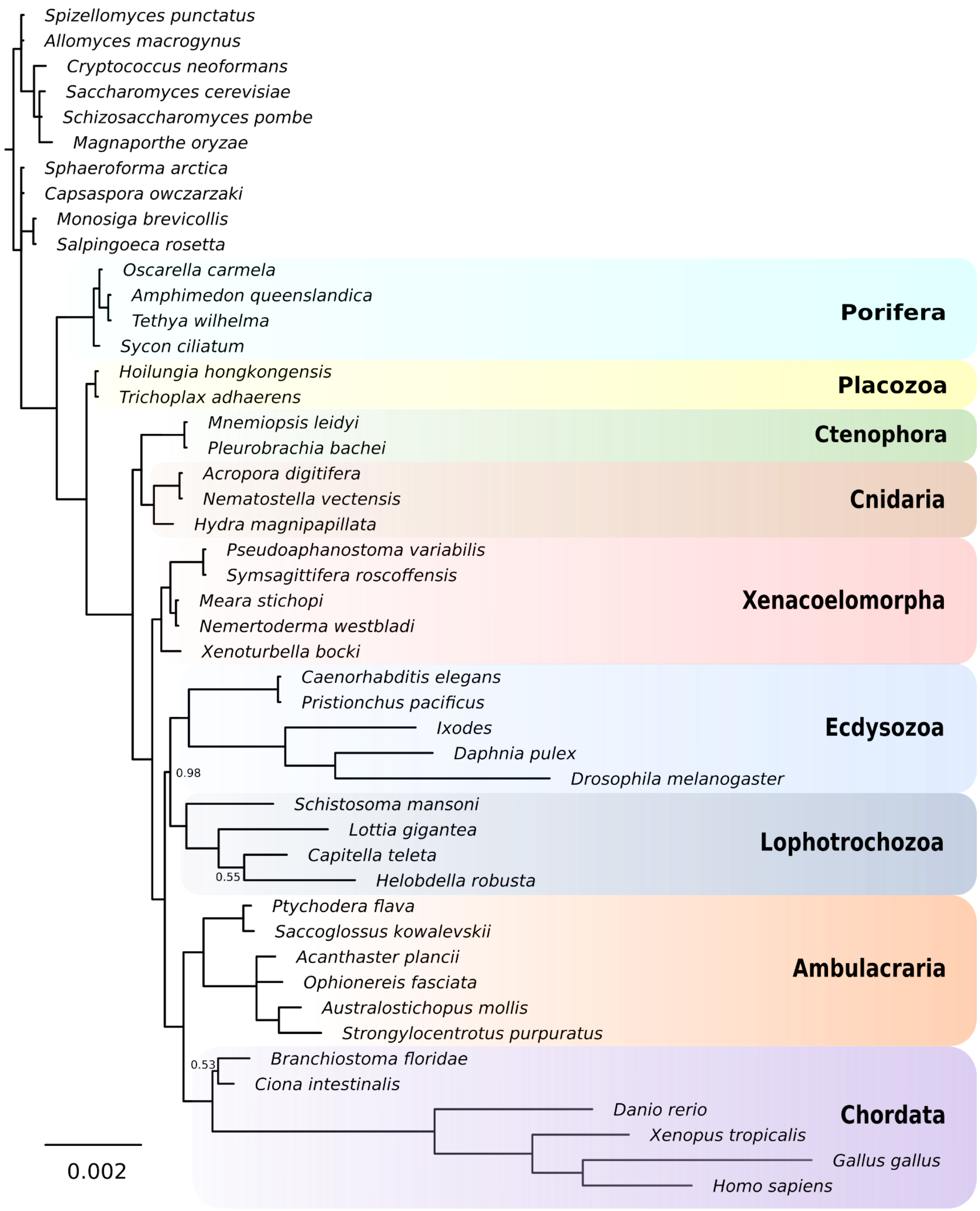


**Supplementary Figure 6: Morphology – non-additive coding, full taxon sample (Bayesian analysis)**. Posterior probabilities lower than 0.99 are indicated.

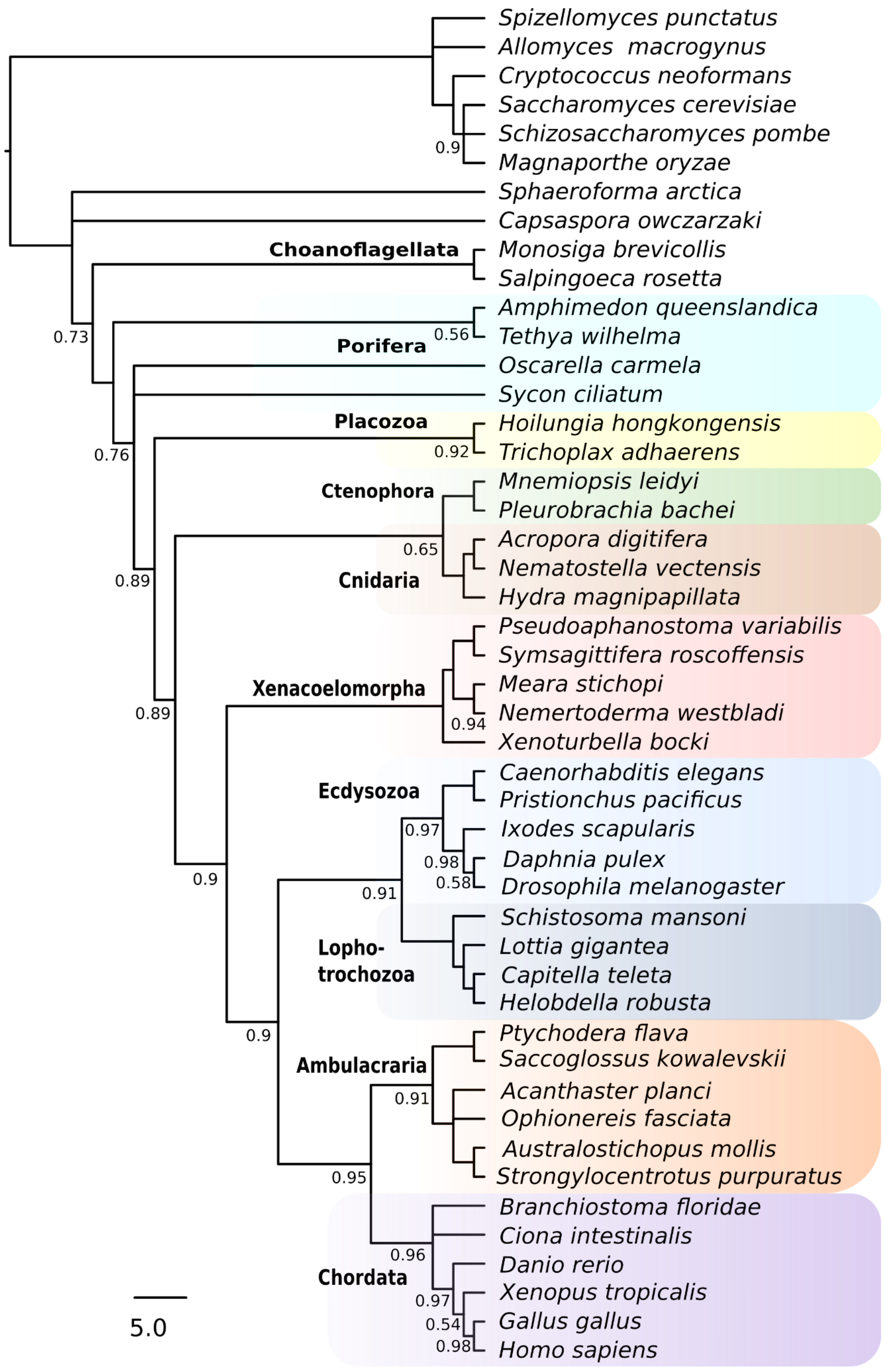


**Supplementary Figure 7: Morphology – reductive coding, full taxon sample (Bayesian analysis).** Posterior probabilities lower than 0.99 are indicated.

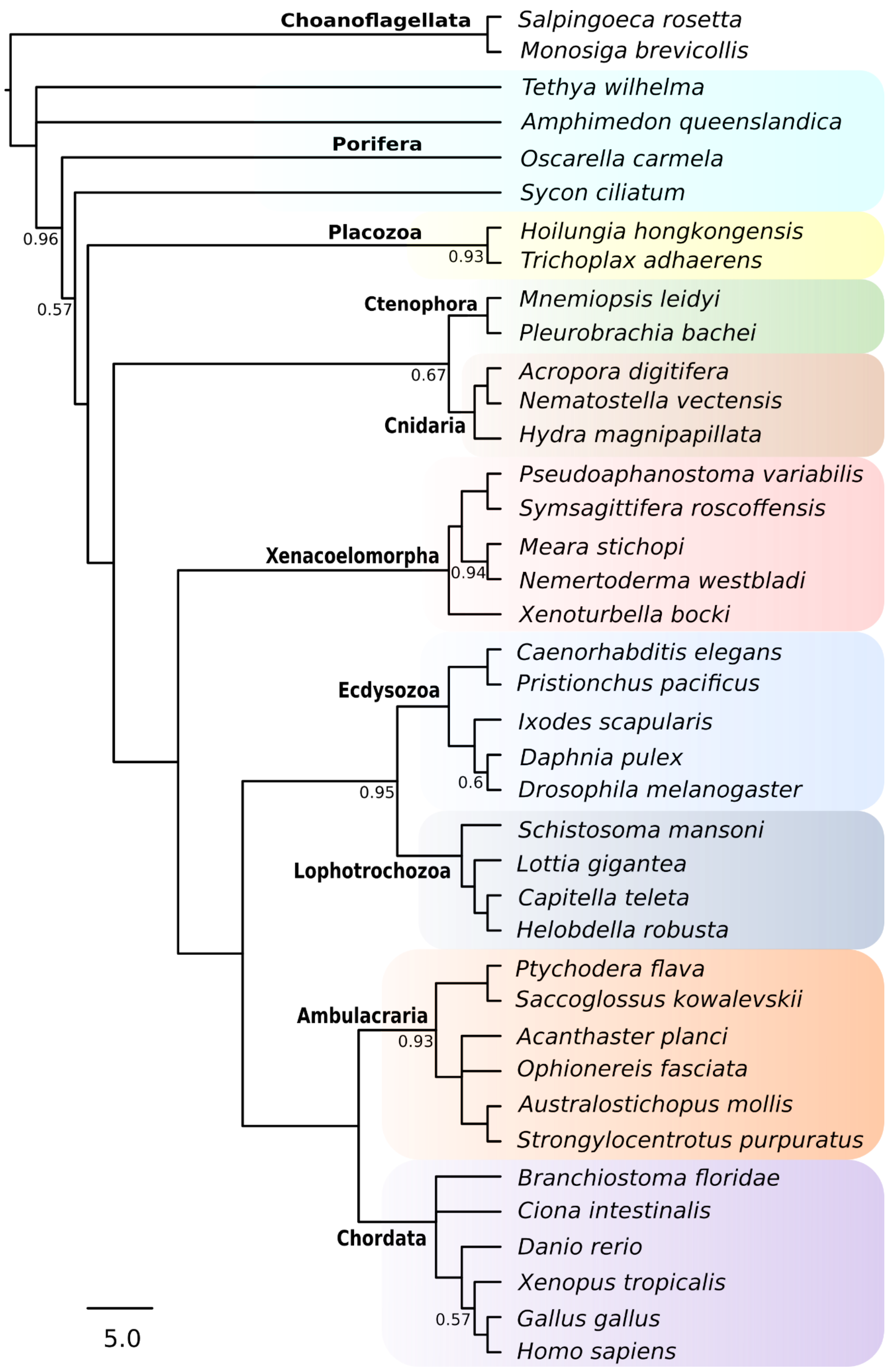


**Supplementary Figure 8: Morphology – reductive coding, reduced outgroup sample (Bayesian analysis).** Posterior probabilities lower than 0.99 are indicated.

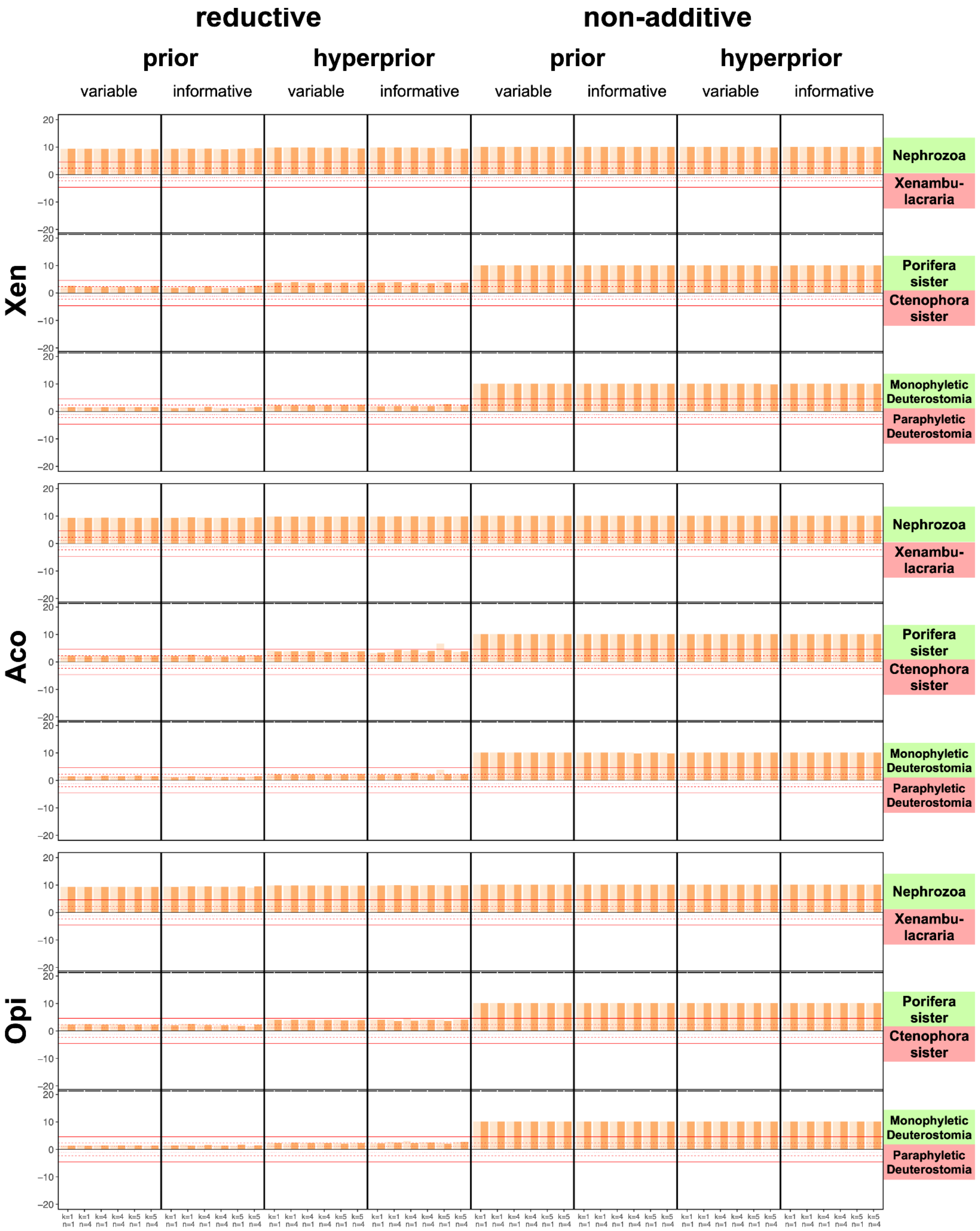


**Supplementary Figure 9: Morphology – Statistical hypothesis testing**. The two different codings are indicated on the top, the three competing hypotheses tested on the right side, and on the left the taxon sampling. Additionally, at the top the different model assumptions about the branch length prior (fixed prior vs hyperprior) and the ascertainment bias correction (invariant vs parsimony informative) are shown. At the bottom the combinations of the different number of rate categories *n* and number of transition rates *k* in the profile mixture are listed. The results of two replicate chains are indicated in shades of orange. For each tested hypothesis, positive values represent support for hypothesis indicated on the right side by light green squares, and negative values represent support for hypothesis indicated on the right side by light red squares. Interpretation of log-posterior odds was done according to Kass and Raftery [13]. Red lines indicate a very strong support level (5,-5).

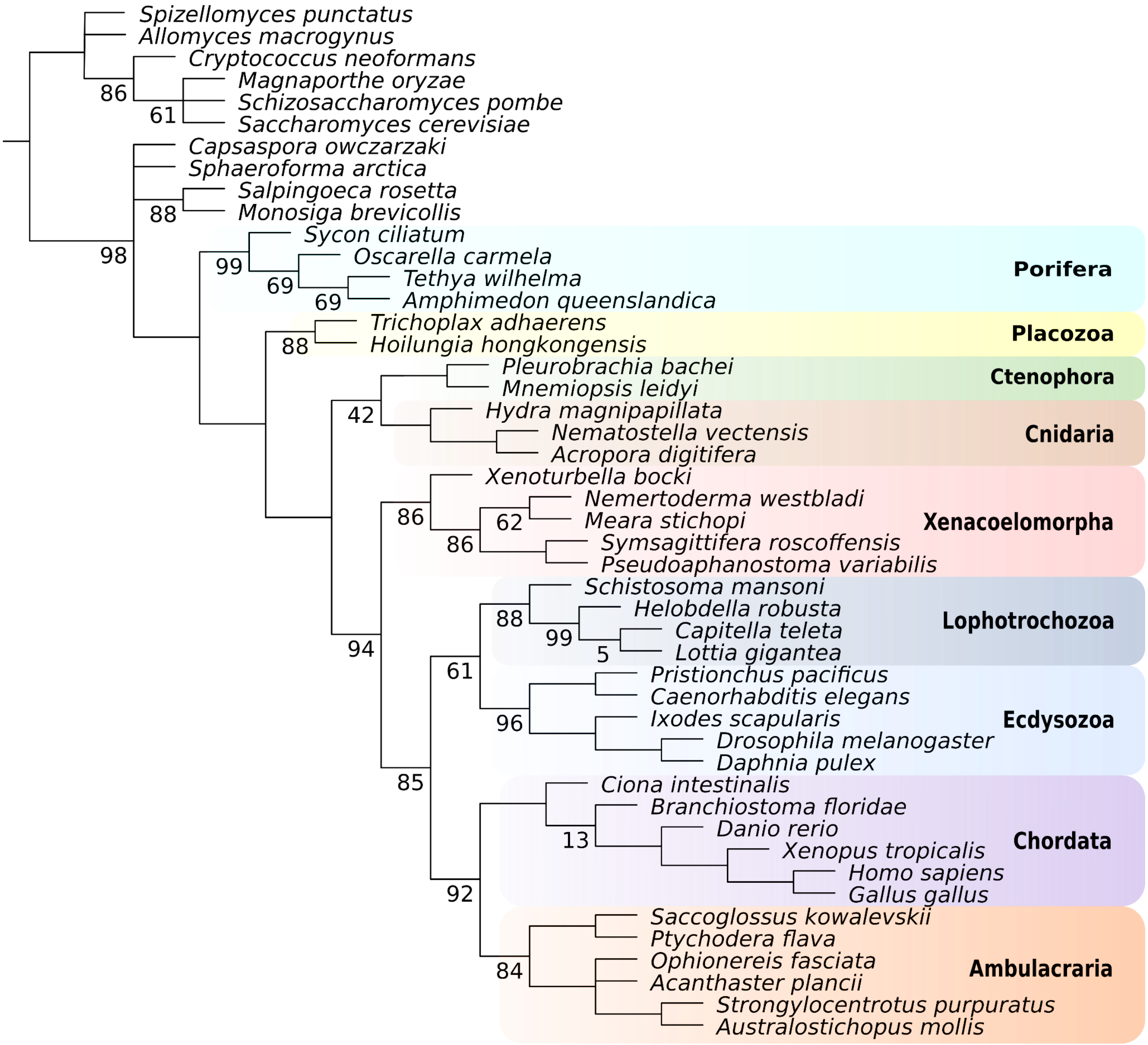


**Supplementary Figure 10: Morphology – non-additive coding, full taxon sample (Maximum Parsimony).** Bootstrap values lower than 100 are indicated.

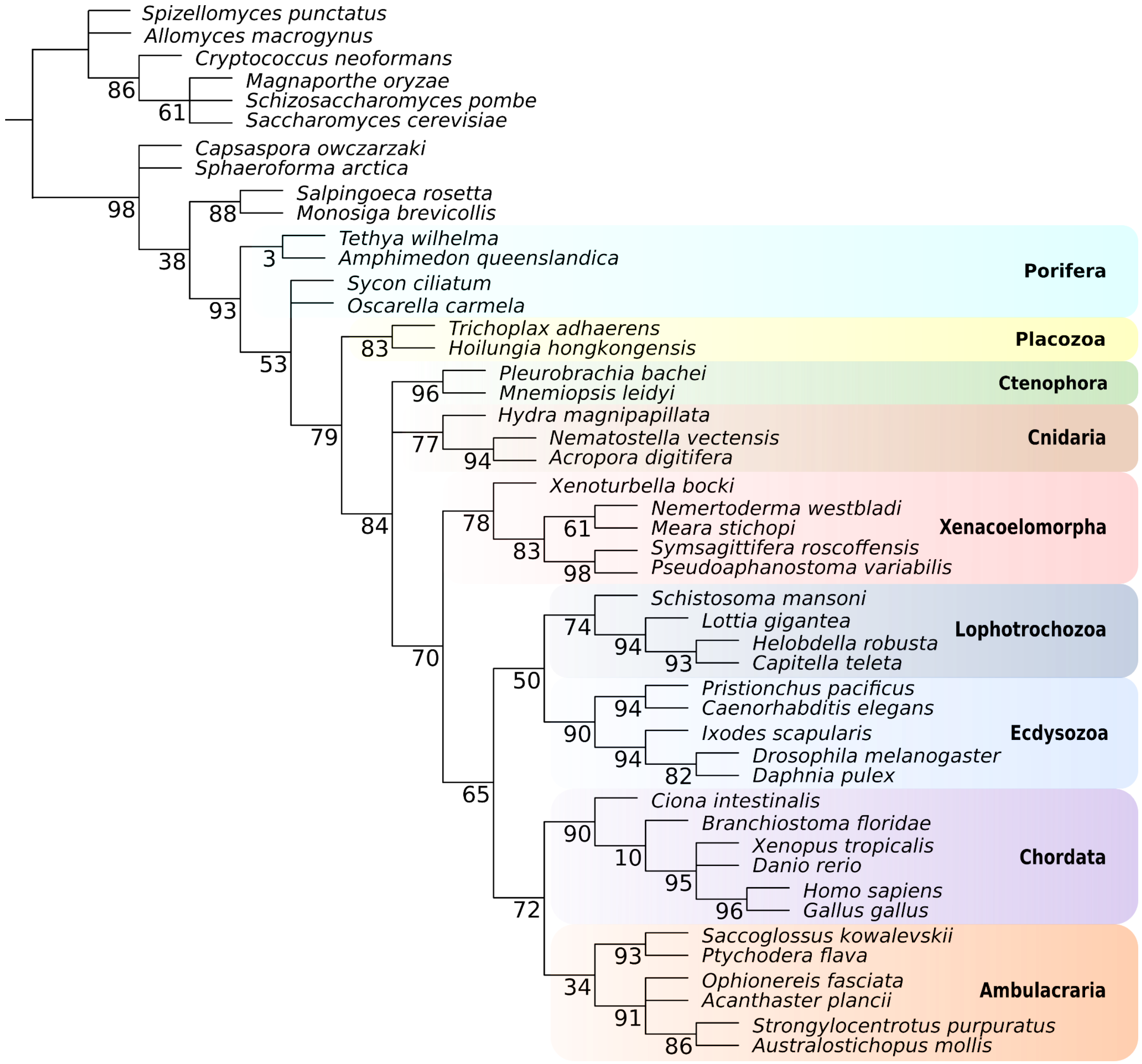


**Supplementary Figure 11: Morphology – reductive coding, full taxon sample (Maximum Parsimony)**. Bootstrap values lower than 100 are indicated.

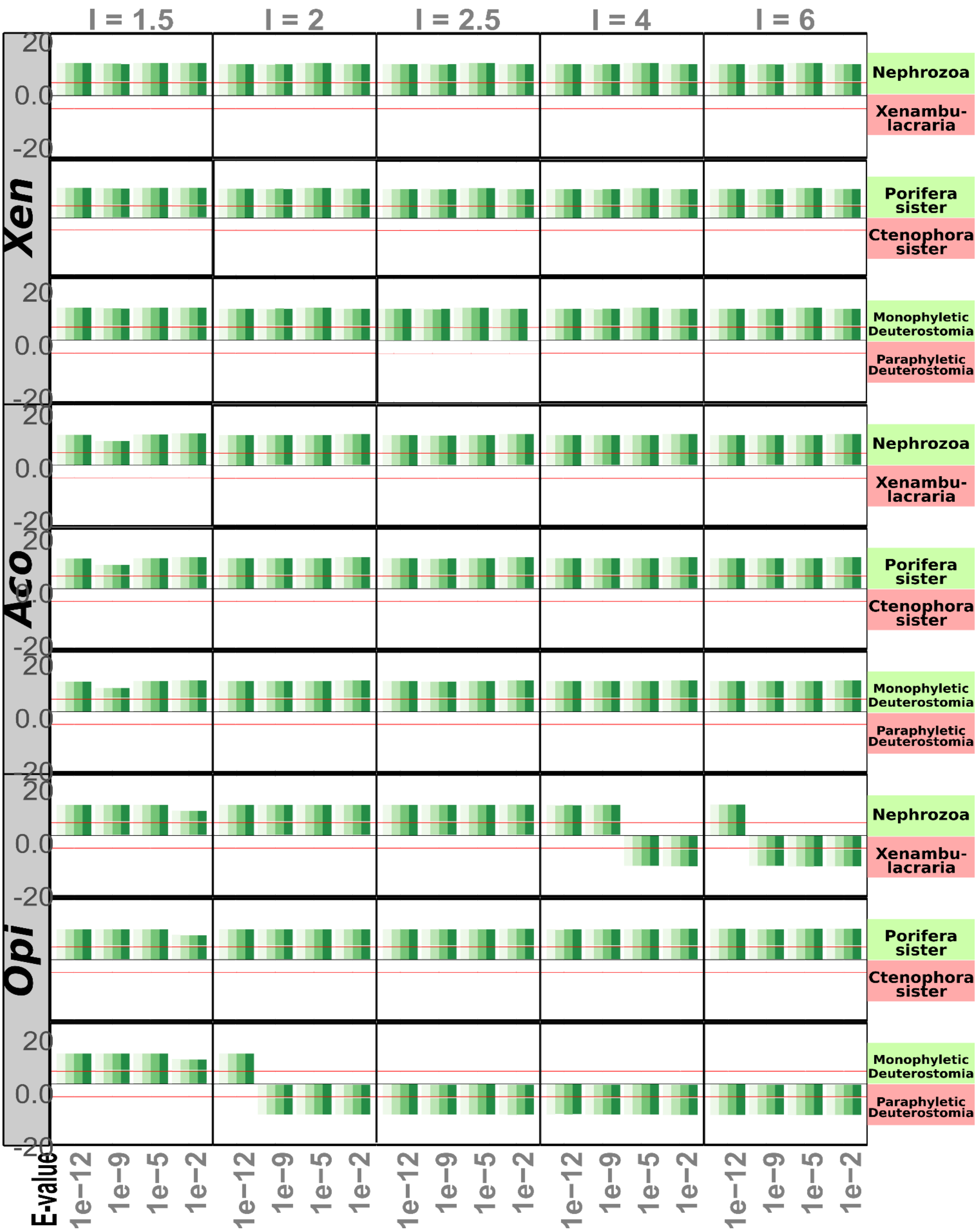


**Supplementary Figure 12: Gene Content – Statistical hypothesis testing for the tested topologies in all orthogroups based datasets.** The three datasets are indicated on the left side, the three competing hypotheses tested are indicated on the right side. The results of the four replicate chains are indicated in shades of green. I-values are indicated on top and E-values on the bottom. For each tested hypothesis, positive values represent support for hypothesis indicated on the right side by light green squares, and negative values represent support for hypothesis indicated on the right side by light red squares Interpretation of log-posterior odds was done according to Kass and Raftery [13]. Red lines indicate a very strong support level (5,-5).

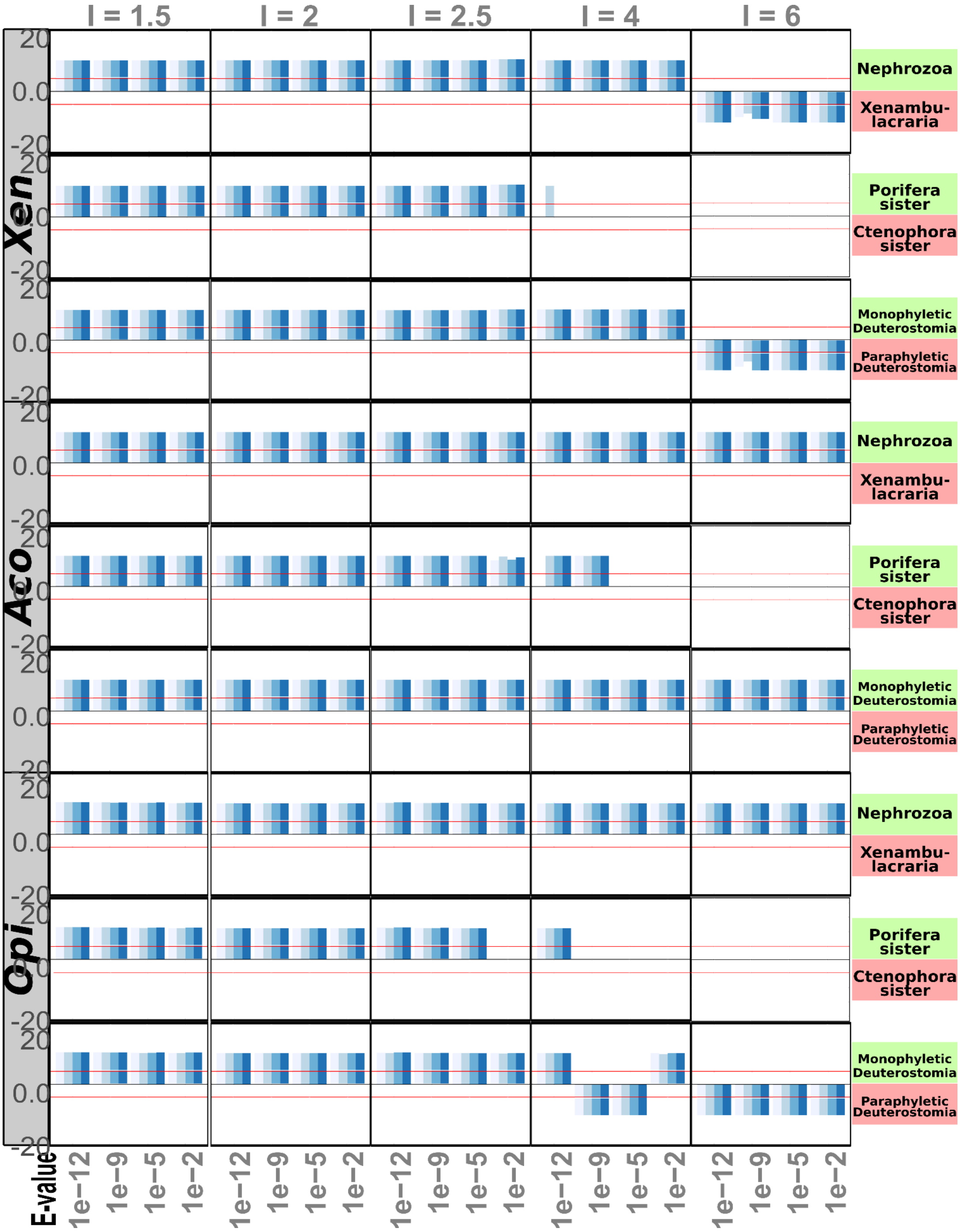


**Supplementary Figure 13: Gene content – Statistical hypothesis testing for the tested topologies in all homogroups based datasets.** The three datasets are indicated on the left side, the three competing hypotheses tested are indicated on the right side. The results of the four replicate chains are indicated in shades of blue. I-values are indicated on top and E-values on the bottom. For each tested hypothesis, positive values represent support for hypothesis indicated on the right side by light green squares, and negative values represent support for hypothesis indicated on the right side by light red squares. No values in the plot mean none of the two tested hypotheses was supported. Interpretation of log-posterior odds was done according to Kass and Raftery [13]. Red lines indicate a very strong support level (5,-5)**.**

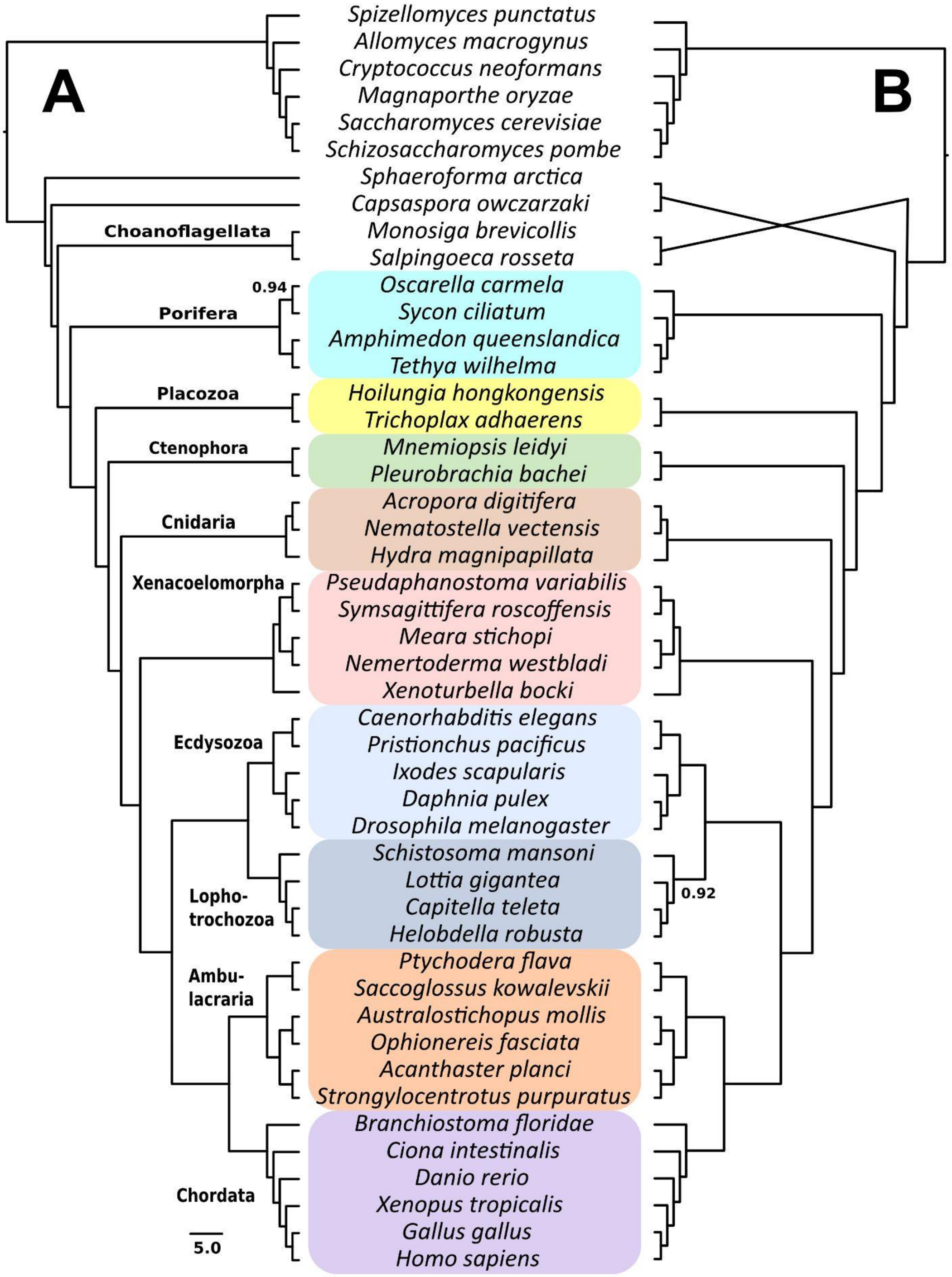


**Supplementary Figure 14: "Total Evidence" Phylogeny of the combined gene content and morphological datasets.** Opi taxon sampling (47 taxa) with the default methods settings for gene content (I-value of 1.5 and E-value of 1e-3) and the reductive coding morphology dataset. A: orthogroups+morphology, B: homogroups+morphology. Posterior probabilities lower than 0.99 are indicated.
